# Matched full-UDG and non-UDG ancient DNA libraries reveal trade-offs in post-mortem damage correction for imputation and kinship inference

**DOI:** 10.64898/2026.09.07.749785

**Authors:** Odongoo Ravdandorj, Chuluun Sampildondov, Khulan Janchiv, Takashi Gakuhari

## Abstract

Ancient DNA studies increasingly combine full uracil-DNA glycosylase-treated, full-UDG, and non-UDG libraries, but which computational damage correction to apply before imputation and kinship analysis remains unsettled. We compared terminal trimming, base-quality rescaling and known-SNP masking in matched full-UDG and non-UDG libraries from the same extracts of two medieval Mongolian individuals. All methods reduced or reversed the difference in mean alternative-allele fraction between damage-prone and transversion SNPs but retained different proportions of covered sites. In non-UDG libraries, masking and rescaling within five bases of each fragment end produced similar cross-library non-reference discordance, NRD, while retaining 92% and 99.3% of covered sites, respectively. In these 3′-biased libraries, ten-base 3′-only trimming retained more covered sites at lower observed discordance than five-base-per- end symmetric trimming. ancIBD inferred widespread sharing of one identity-by-descent chromosome copy, IBD1, across correction methods. Uncorrected non-UDG data shifted TKGWV2 toward second-degree estimates, whereas corrected mixed-library comparisons supported first-degree relatedness. READv2 and the low rate of opposite-homozygote sites, IBS0, supported a parent-offspring relationship. Mitochondrial data and genetic sex favoured ORT16 as mother and ORT15 as son. No method performed best across all measures, so the choice depends on whether residual damage, site retention or the downstream analysis matters most.

## Introduction

Ancient DNA (aDNA) molecules are typically short, chemically modified, and present in small amounts. After death, hydrolytic depurination removes adenine or guanine bases, leaving abasic sites that make the DNA strand susceptible to breakage. The resulting fragments often contain short single-stranded overhangs. Cytosine deamination occurs at a higher rate in these overhangs than in double-stranded DNA and converts cytosine to uracil.[1] When these deaminated cytosines (uracils) are copied by a polymerase capable of reading through uracil, the polymerase inserts adenine opposite it, causing the original cytosine to be represented as thymine in the sequence data. In double-stranded libraries, this damage appears as C→T substitutions near the 5′ ends of sequencing reads and complementary G→A substitutions near the 3′ ends.[1,2] In single-stranded libraries, each original DNA strand is converted independently, and C→T substitutions are therefore observed at both ends.[3,4] These terminal patterns are widely used to authenticate ancient DNA, but the same damage can be mistaken for genuine genetic variation and introduce errors into genotype calling, imputation, and kinship analyses.

The effects of post-mortem cytosine deamination can be reduced enzymatically before sequencing or addressed computationally afterward. Two common enzymatic approaches are full-UDG treatment and partial-UDG treatment, also called half UDG.[5] The uracil-specific excision reagent (USER) used in these treatments contains uracil-DNA glycosylase (UDG) and endonuclease VIII. UDG recognises and excises uracil bases, leaving abasic sites.

Endonuclease VIII then cleaves the DNA backbone at these sites.[2] In full-UDG treatment, DNA is incubated for an extended period with USER and T4 polynucleotide kinase (T4 PNK), without adding UDG inhibitor (UGI).

T4 PNK prepares the terminal phosphate groups, and T4 DNA polymerase is then added to fill or trim uneven DNA ends. This treatment removes most uracils throughout the DNA fragments, including near their ends. It therefore reduces uracils but also removes most of the terminal signal used to authenticate ancient DNA.[2,5] In partial-UDG treatment, USER acts for a shorter period, after which UDG is inhibited with UGI before T4 PNK and T4 DNA polymerase are added.[5] This removes most internal uracils while retaining part of the terminal damage signal. Consequently, a few terminal bases are often excluded from partial-UDG libraries before downstream analysis.[6] Residual terminal damage can also persist even after full-UDG treatment, so the observed damage profile should determine whether terminal trimming is necessary.[2]

Many published aDNA datasets were generated without UDG treatment (hereafter, non-UDG) and therefore retain cytosine deamination damage throughout the DNA fragments, with the highest frequencies near their ends.[1] When non-UDG data are analysed together with full-UDG or partial-UDG data, differences in residual damage may produce different genotyping error rates and affect imputation accuracy.[6,7] Unequal deamination may also bias pairwise mismatch statistics used by kinship methods such as READv2.[8] Several computational approaches are available to reduce the influence of post-mortem damage (PMD). Here, we used trimBam in bamUtil to soft-clip a fixed number of terminal bases, retaining their sequence in the BAM but excluding them from genotype likelihoods.[6,9] mapDamage2 estimates position-specific damage probabilities and rescales base quality scores accordingly.[10] bamRefine uses a predefined SNP catalogue to mask observations at PMD-susceptible sites within a selected terminal window.[6]

Previous work compared terminal trimming, mapDamage2 rescaling and bamRefine masking in simulated paleogenomes, and trimming with bamRefine in empirical paleogenomes, focusing on genotyping error, reference bias and data loss.[6] Low-coverage ancient genomes contain too few observations to determine many genotypes directly. Imputation estimates missing genotypes by comparing the observed data with haplotypes in a reference panel; here, we used GLIMPSE with the 1000 Genomes Project panel.[11–13] The resulting imputed data can then be analysed with tools like ancIBD to identify identity-by-descent (IBD) segments, chromosomal regions that two individuals inherited from a recent common ancestor.[14] Here, we extend the earlier comparison to full-UDG and non-UDG libraries prepared from the same individuals and test whether processing choices affect genotype agreement and kinship estimates.

We studied two medieval Mongolian individuals, ORT15 and ORT16, selected because preliminary analysis of the screening data suggested a first-degree relationship. Because full-UDG treatment removes most of the terminal signal used for authentication, we first prepared one non-UDG library to assess ancient DNA authenticity and endogenous DNA content. This library received no enzymatic damage removal before library preparation, but was exposed to USER during loop-adaptor opening after ligation, which affects the terminal damage profile. We subsequently prepared three full-UDG libraries from separate aliquots of the same source extracts. The library types were therefore matched by individual and source extract, but not by sequencing depth. We analysed the full-UDG and non-UDG datasets separately using trimBam, mapDamage2, and bamRefine. We tested different amounts of correction near the fragment ends, including longer ranges for the more damaged non-UDG libraries. We first examined how much damage remained and how many SNPs were still available after each correction. We then imputed each dataset separately and compared the genotypes obtained from the two library types for the same individual. Finally, we tested whether the correction method or the combination of library types changed the inferred relationship between ORT15 and ORT16. We compared results obtained from two full-UDG datasets, two non-UDG datasets, and one dataset of each type. Because the full-UDG data were also low coverage and their missing genotypes had to be inferred, they could not provide a set of known correct genotypes. Instead, they provided a lower-damage comparison. Our study therefore evaluates the consistency and practical limitations of the three methods rather than determining absolute genotype accuracy. Figure 1 summarises the study design.

**Figure 1.**
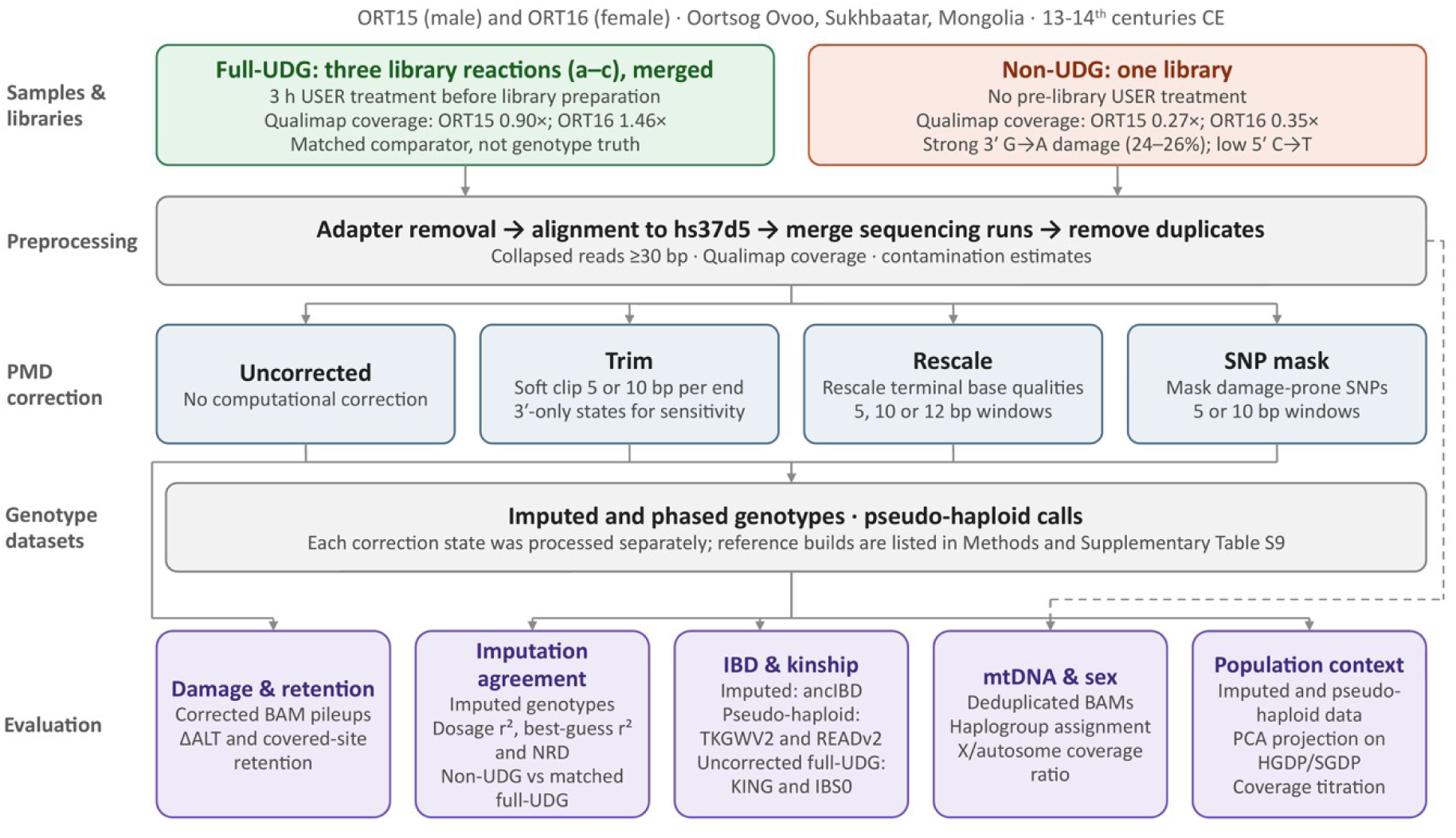
Overview of the study workflow. ORT15 and ORT16 each had three full-UDG library reactions, merged for analysis, and one lower-coverage non-UDG library. Rounded Qualimap coverage was 0.90× for ORT15 full-UDG, 0.27× for ORT15 non-UDG, 1.46× for ORT16 full-UDG and 0.35× for ORT16 non-UDG. Following shared preprocessing, the data were analysed without computational correction and after terminal trimming, base-quality rescaling or SNP masking. The dashed arrow shows that mitochondrial haplogroup assignment and genetic-sex estimation used deduplicated BAMs directly, bypassing computational damage correction and genotype generation. Damage-associated allele support and covered-site retention were assessed within correction states; imputation agreement compared each non-UDG state with the matched full-UDG data from the same individual; IBD and kinship compared ORT15 with ORT16; and PCA assessed broad population position. Full-UDG data served as matched comparators, not known genotype truth.

## Results

### Library quality comparison between library types

Full-UDG libraries showed low terminal damage, whereas the non-UDG libraries showed a strong G→A signal at the 3′ end (Supplementary Fig. S1; Supplementary Table S1). At the terminal 3′ position, G→A frequencies were 1.30% in ORT15 and 1.34% in ORT16 full-UDG data, compared with 26.02% and 23.92%, respectively, in the non-UDG data. By contrast, at the terminal 5′ position, C→T frequencies in the non-UDG libraries were much lower, at 2.60% and 3.59%, respectively. The difference between the two ends may reflect the USER step used to open the loop adaptor after ligation. Here, non-UDG means that no enzymatic damage-removal treatment was performed before library preparation, although the libraries were later exposed to USER (see Methods).

Contamination estimates were low (Supplementary Table S2). In the genetically male individual ORT15, ANGSD Method 1 estimated present-day human nuclear contamination from 6757 covered HapMapChrX sites. The estimate was 0.22% by method of moments (SE, 0.08%) and 0.39% by maximum likelihood (SE, 0.12%). This X-chromosome method was not applicable to genetically female ORT16. Standalone mtCont estimated mitochondrial contamination at 1% in each individual, with a 95% confidence interval of 0–2% for both. Overall, these results indicated low mitochondrial contamination in both individuals and low nuclear contamination in ORT15, supporting the use of these libraries in downstream analysis.

In the merged full-UDG datasets, adding other read categories provided little additional mean genomic coverage. Including truncated-collapsed reads increased coverage by 0.004× in ORT15 and 0.0049× in ORT16, while the endogenous DNA proportion changed by less than 0.5 percentage points in either individual. Including unmerged paired-end reads resulted in further increases of only 0.0019× and 0.0024×, respectively, but reduced the endogenous DNA proportion from 66.75% to 58.61% in ORT15 and from 82.76% to 76.17% in ORT16. Based on these small coverage gains, we used collapsed reads alone for all subsequent imputation and IBD analyses.

### PMD correction changed the damage-associated ALT-fraction contrast and covered-site retention

Because cytosine deamination can increase apparent ALT support at C→T/G→A SNPs, we used the study-specific ΔALT contrast defined in Methods. Values are reported in percentage points. Positive values indicate greater ALT support at these damage-prone sites and are compatible with residual deamination, although ΔALT is not a direct estimate of PMD. Before correction, ΔALT was higher in the non-UDG data than in the matched full-UDG data for both individuals (ORT15, 0.85 versus 0.47 percentage points; ORT16, 0.64 versus 0.34; Fig. 2; Supplementary Table S3). In the primary analysis, which used the SNPs eligible within each library, ΔALT was 0.38 percentage points higher in non-UDG than in full-UDG data for ORT15 (95% CI, 0.29–0.48) and 0.30 percentage points higher for ORT16 (95% CI, 0.19–0.41). These intervals were obtained using a paired jackknife that omitted one autosome at a time.

**Figure 2.**
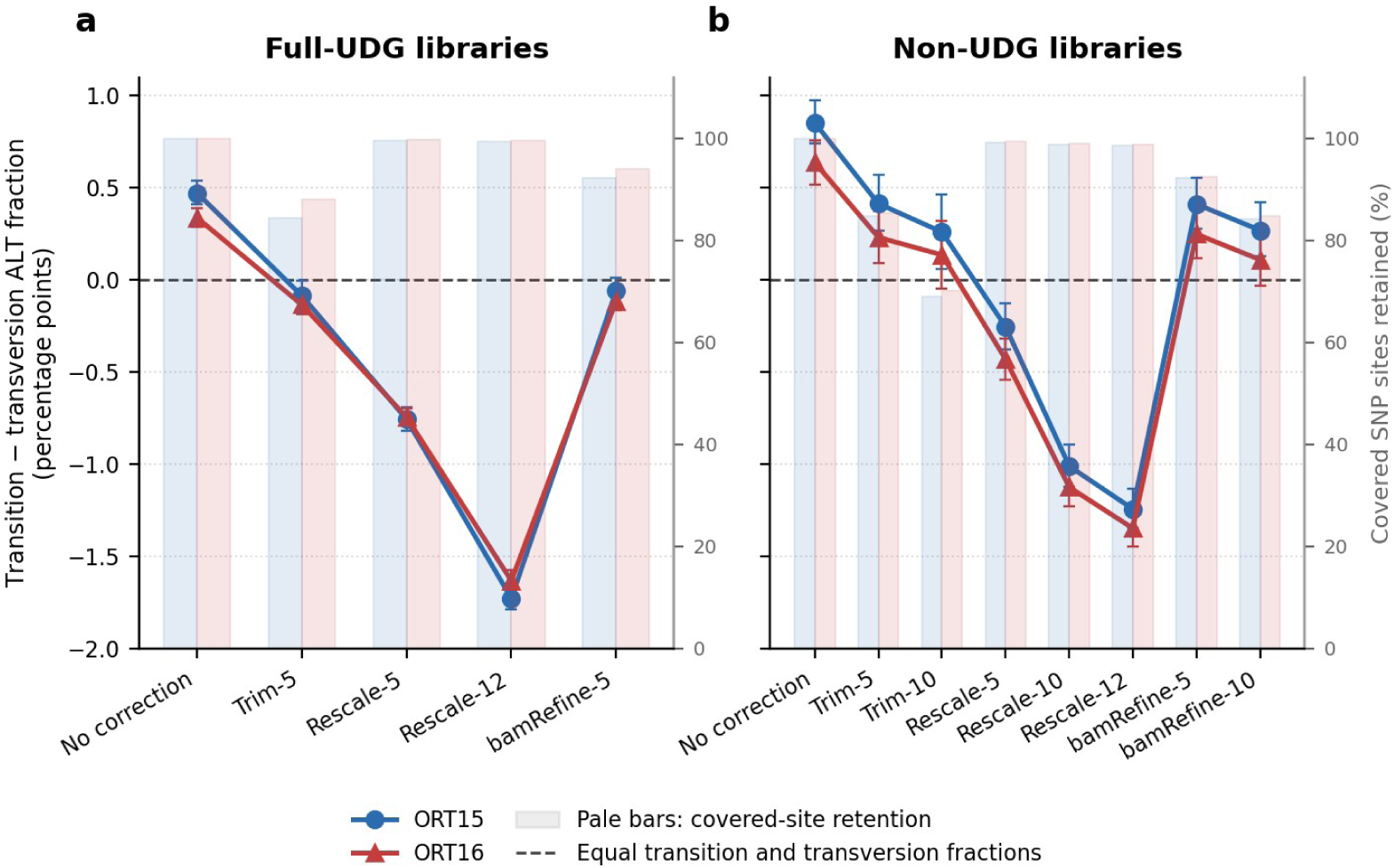
Damage-associated allele support and covered-site retention after PMD correction. Panel **(a)** shows full-UDG libraries and panel **(b)** shows non-UDG libraries. Lines show ΔALT, defined as the mean alternative-allele fraction at C→T/G→A SNPs minus the mean at transversion SNPs; these means require combined REF+ALT depth ≥ 3. Positive values indicate greater alternative-allele support at damage-prone sites. Pale bars show the percentage of the uncorrected covered SNP set retained after correction, requiring combined REF+ALT depth ≥ 1. Error bars are leave-one-autosome-out 95% confidence intervals for ΔALT; retention is shown descriptively without intervals. Because SNP composition and eligible covered sites can differ across classes and correction states, ΔALT is descriptive; a negative ΔALT alone does not demonstrate loss of genuine variants. Values are reported in Supplementary Table S3. The two 3′-only sensitivity states are included in that table but are not plotted.

All correction methods lowered ΔALT in the non-UDG libraries, but the pattern differed by method. Trim-5 and bamRefine-5 left small positive contrasts, whereas mapDamage2 rescaling shifted ΔALT below zero in both non-UDG and full-UDG libraries. The downward shift was larger with the 12-base window than with the 5-base window. Under the primary autosome jackknife, the 95% CIs for three individual ΔALT estimates included zero: ORT15 full-UDG bamRefine-5 and ORT16 non-UDG Trim-10 and bamRefine-10 (Supplementary Table S3). For all three estimates, both the 5-Mb and 10-Mb block intervals excluded zero, whereas the primary autosome-jackknife intervals included zero. In contrast, every paired 95% CI comparing a corrected dataset with its uncorrected counterpart lay entirely below zero under the autosome, 5-Mb and 10-Mb analyses.

When we repeated each comparison using only SNPs eligible in all contributing datasets, several estimated differences changed in magnitude. Under this restriction, the interval for the difference between uncorrected non-UDG and full-UDG data in ORT15 included zero. For ORT16, the point estimate comparing the Rescale-12 change in non-UDG data with the corresponding change in full-UDG data changed sign relative to the primary analysis (Supplementary Data). Because the primary estimate was close to zero and its confidence interval included zero, this was a sign change in the estimate rather than evidence that the underlying effect reversed.

Because transition and transversion SNP classes may differ in their allele-frequency distributions and REF/ALT composition, and because each processed dataset retained a different set of eligible SNPs, ΔALT depends partly on which SNPs are analysed and should be interpreted descriptively. mapDamage2 fitted its damage model and rescaled base qualities independently for each library, so the same window length did not necessarily produce identical adjustments. Negative ΔALT values alone do not demonstrate overcorrection or loss of genuine variants.

To assess the loss of covered SNPs, we counted SNPs with at least one read that passed the mapping-quality and base-quality filters and supported either REF or ALT. We expressed the corrected count as a percentage of the corresponding count in the uncorrected library. At the five-base setting shared by all three methods, Rescale-5 retained 99.3% of covered SNPs in both individuals, bamRefine-5 retained 92.2–92.4%, and Trim-5 retained 84.7–85.3%. More extensive trimming with Trim-10 reduced retention to 69.0% in ORT15 and 70.1% in ORT16. Rescale-5 therefore retained the highest proportion of covered SNPs among the tested corrections, although it also shifted ΔALT below zero (Fig. 2; Supplementary Table S3).

Covered-site retention indicates only whether at least one qualifying read remained at a SNP; it does not measure retained read depth or genotype accuracy.

An additional end-specific sensitivity analysis evaluated clipping confined to the 3′ end of the non-UDG reads (Table 1; Supplementary Table S3). In the prespecified window-matched comparison, 3′-only-5 retained 93.4% and 93.9% of covered SNPs in ORT15 and ORT16, respectively, compared with 84.7% and 85.3% after symmetric Trim-5. The corresponding NRD values were 18.60% and 15.02% after 3′-only-5, versus 19.42% and 15.69% after symmetric Trim-5. However, 3′-only-5 left higher ΔALT values: the paired leave-one-autosome-out differences, calculated as 3′-only-5 minus symmetric Trim-5, were 0.133 percentage points in ORT15 (95% CI, 0.074–0.192) and 0.088 in ORT16 (0.058–0.119). In the equal-total-bases comparison, 3′-only-10 and symmetric Trim-5 each clipped ten bases per read. The 3′-only-10 state retained 86.9% and 87.8% of covered SNPs and gave NRD values of 18.82% and 15.29%. Its paired ΔALT differences from symmetric Trim-5 were 0.046 percentage points in ORT15 (95% CI, −0.046 to 0.138) and 0.008 in ORT16 (−0.046 to 0.062). The 5-Mb and 10-Mb block jackknives supported the same inferences.

**Table 1.** Cross-library imputation agreement before and after PMD correction. Each uncorrected or corrected non-UDG dataset was evaluated against the same uncorrected, imputed full-UDG comparator from that individual. The 3′-only-5 and 3′-only-10 states soft-clipped 5 or 10 bases, respectively, from the strand-aware 3′ end only, whereas Trim-5 and Trim-10 soft-clipped the stated number from both ends. Dosage and best-guess *r*² are count-weighted means for variants with MAF≥0.1%; NRD includes all evaluable non-reference comparisons. Dosage *r*² uses expected alternate-allele dosage derived from posterior genotype probabilities, whereas best-guess *r*² uses the most probable genotype.

| Sample | Strategy | Dosage $r^2$ | Best-guess $r^2$ | NRD (%) |
| --- | --- | --- | --- | --- |
| ORT15 | Uncorrected | 0.4354 | 0.4175 | 19.23 |
| ORT15 | Trim-5 | 0.4474 | 0.4258 | 19.42 |
| ORT15 | Trim-10 | 0.4364 | 0.4111 | 21.33 |
| ORT15 | 3'-only-5 | 0.4545 | 0.4343 | 18.60 |
| ORT15 | 3'-only-10 | 0.4582 | 0.4359 | 18.82 |
| ORT15 | Rescale-5 | 0.4630 | 0.4413 | 18.32 |
| ORT15 | Rescale-10 | 0.4698 | 0.4456 | 18.37 |
| ORT15 | Rescale-12 | 0.4699 | 0.4448 | 18.61 |
| ORT15 | bamRefine-5 | 0.4622 | 0.4406 | 18.30 |
| ORT15 | bamRefine-10 | 0.4664 | 0.4434 | 18.57 |
| ORT16 | Uncorrected | 0.4873 | 0.4698 | 15.60 |
| ORT16 | Trim-5 | 0.5041 | 0.4813 | 15.69 |
| ORT16 | Trim-10 | 0.4898 | 0.4624 | 17.51 |
| ORT16 | 3'-only-5 | 0.5099 | 0.4902 | 15.02 |
| ORT16 | 3'-only-10 | 0.5103 | 0.4871 | 15.29 |
| ORT16 | Rescale-5 | 0.5159 | 0.4933 | 14.85 |
| ORT16 | Rescale-10 | 0.5203 | 0.4943 | 15.05 |
| ORT16 | Rescale-12 | 0.5205 | 0.4944 | 15.15 |
| ORT16 | bamRefine-5 | 0.5156 | 0.4945 | 14.88 |
| ORT16 | bamRefine-10 | 0.5156 | 0.4897 | 15.36 |

*Note: these measures quantify agreement between library types, not accuracy against known genotypes. NRD was calculated independently for each strategy over its method-specific evaluable site set; comparisons among NRD values are therefore descriptive rather than site matched.*

Read-end C→T and G→A mismatch frequencies, calculated with mapDamage2, showed a clear reduction after terminal soft-clipping and only minor changes after bamRefine masking. In Supplementary Fig. S2, Rescale-5 and the uncorrected data have the same curve because rescaling changes base-quality scores rather than the recorded bases, and the profiles were calculated with a minimum base-quality threshold of zero. These profiles illustrate how soft-clipping and masking alter recorded read-end mismatches but do not provide a comparable measure of downstream effects across all three methods.

Together, ΔALT and covered-site retention described changes in damage-associated ALT support and SNP availability, but neither measured genotype accuracy. We therefore evaluated practical downstream effects using cross-library genotype agreement and kinship analyses.

### Rescaling and masking showed the most consistent increases in cross-library agreement

For each individual, we compared imputed genotypes from each non-UDG correction state with the same uncorrected, independently imputed full-UDG dataset. Higher dosage and best-guess r² values and lower non-reference discordance (NRD) indicate closer agreement between library types. Rescaling and bamRefine masking numerically improved all three measures relative to the uncorrected non-UDG data (Fig. 3; Table 1). In ORT15, NRD decreased from 19.23% to 18.30% with bamRefine-5 and 18.32% with Rescale-5. In ORT16, it decreased from 15.60% to 14.88% and 14.85%, respectively. Trim-5 slightly increased both r² measures but also increased NRD to 19.42% and 15.69%. Trim-10 increased NRD more clearly, to 21.33% and 17.51%, and reduced best-guess r² below the uncorrected value, while dosage r² remained nearly unchanged. Thus, both trimming settings increased NRD relative to the uncorrected data, whereas every rescaling and masking setting reduced it; effects on the r² measures were smaller and did not always follow the same pattern. Because the full-UDG comparator was also low-coverage and imputed, and because the library types differed in sequencing depth, these changes measure cross-library agreement rather than genotype accuracy.

**Figure 3.**
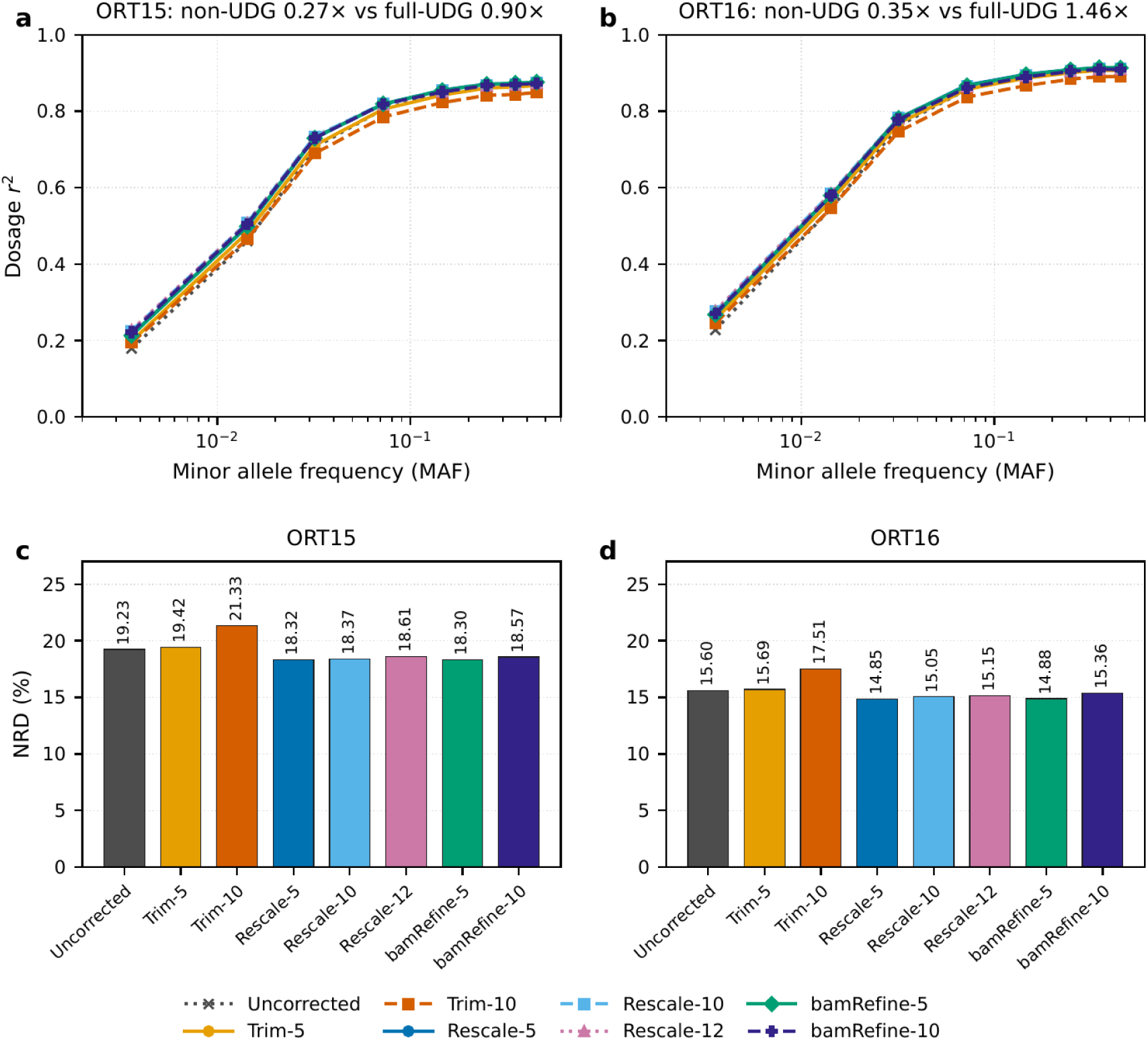
Cross-library imputation agreement before and after PMD correction. Panels **(a)** and **(b)** show dosage r² by minor allele frequency bin (MAF ≥ 0.1%) for the eight primary non-UDG states from ORT15 (0.27×) and ORT16 (0.35×), respectively. Each state is compared with the same uncorrected full-UDG dataset from that individual (0.90× and 1.46×, respectively). These are rounded Qualimap coverage estimates. Panels **(c)** and **(d)** show non-reference discordance (NRD) across all evaluable non-reference genotype comparisons; axes begin at zero. Higher r² and lower NRD indicate closer agreement between library types, not accuracy against known genotypes. Best-guess r² and exact values for these eight states, plus the two 3′-only trimming sensitivity states not shown here, are reported in Table 1.

Dosage r² was lower for rare variants and generally increased with minor allele frequency (MAF). In both individuals, the uncorrected data had the lowest dosage r² in the two lowest plotted bins, 0.1–1% and 1–2%, whereas Trim-10 had the lowest dosage r² in every bin from 2% upward. No uncertainty intervals were calculated for individual MAF bins, so these rankings are descriptive.

No method ranked first across all measures. Rescale-12 had the highest count-weighted dosage r² in both individuals, whereas the highest best-guess r² occurred with Rescale-10 in ORT15 and with bamRefine-5 in ORT16. The minimum NRD occurred with bamRefine-5 in ORT15 and with Rescale-5 in ORT16. The NRD difference between these two five-base methods was only 0.02–0.03 percentage points. Paired delete-one-autosome jackknife 95% confidence intervals included zero in both individuals (ORT15, −0.203 to 0.161 percentage points; ORT16, −0.107 to 0.168; Supplementary Table S4). Thus, no NRD difference was detected in either individual, although equivalence was not established. These intervals describe variation among autosomes within each individual, not replication across individuals. Because the observed NRD values were nearly identical, we next assessed whether the first-degree IBD signal remained stable despite the methods’ different covered-site retention.

### IBD1 sharing persisted across PMD correction methods

We first kept ORT15’s full-UDG data uncorrected while varying the correction applied to ORT16’s non-UDG data. We then repeated the comparison in the opposite direction, keeping ORT16’s full-UDG data uncorrected while varying the correction applied to ORT15’s non-UDG data. Rescale-5 produced the largest observed total IBD1 length in both directions: 3027.9 cM for ORT15 full-UDG × ORT16 non-UDG and 2880.5 cM for ORT15 non-UDG × ORT16 full-UDG (Fig. 4a,b; Supplementary Table S5).

**Figure 4.**
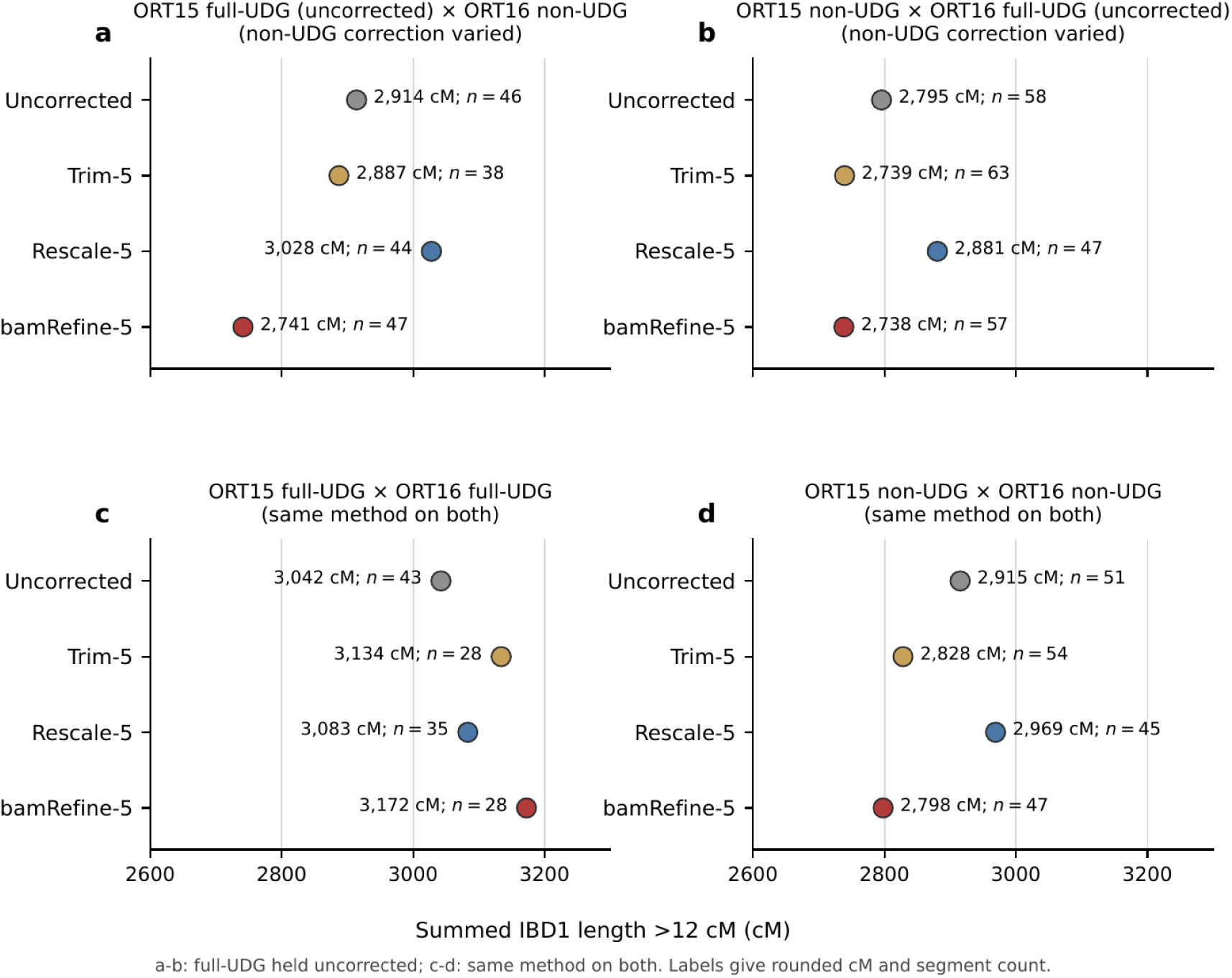
IBD1 length in asymmetric and same-method comparisons. Points show the summed length of IBD1 segments longer than 12 cM with marker density above 220 SNPs/cM; labels show the rounded total and number of retained segments. The x axis is truncated to 2600–3300 cM in every panel. In panel **(a)**, ORT15 full-UDG remains uncorrected while correction of ORT16 non-UDG varies; panel **(b)** reverses these roles. In panels **(c)** and **(d)**, the same state is applied to both full-UDG libraries or both non-UDG libraries, respectively. Because both members change together in panels (c) and (d), differences cannot be attributed to one corrected dataset. Without known truth, a larger IBD1 total shows only that more sharing was inferred under that processing state, not that it was more accurate. The full matrix and a diagnostic relating total length to segment count are shown in Supplementary Figures S7 and S8.

We next compared the two individuals using the same library type on both sides: full-UDG × full-UDG and non-UDG × non-UDG. Each pairing was evaluated with both datasets uncorrected and with both processed using Trim-5, Rescale-5 or bamRefine-5. Among the full-UDG comparisons, bamRefine-5 produced the largest observed total IBD1 length (3172.5 cM).

Among the non-UDG comparisons, Rescale-5 produced the largest observed total IBD1 length (2969.0 cM; Fig. 4c,d). Because both datasets changed between processing states, these comparisons could not isolate the effect of changing only one dataset.

Finally, we returned to the mixed-library pairings, using full-UDG data for one individual and non-UDG data for the other, and applied the same processing state to both inputs. Each of the four states (uncorrected, Trim-5, Rescale-5, and bamRefine-5) was examined in both library directions, giving eight comparisons. Filtered IBD1 segments were detected on 20 or 21 of the 22 autosomes, depending on the processing state and library direction (Fig. 5). Summed IBD1 ranged from 2785.9 to 2988.3 cM, equivalent to 78.7–84.4% of the autosomal genetic-map span defined by the common marker panel (3538.85 cM). The different designs explain why Rescale-5 yielded 3027.9 cM in Fig. 4a, where only the non-UDG input was rescaled, but 2988.3 cM in Fig. 5, where both inputs were rescaled.

**Figure 5.**
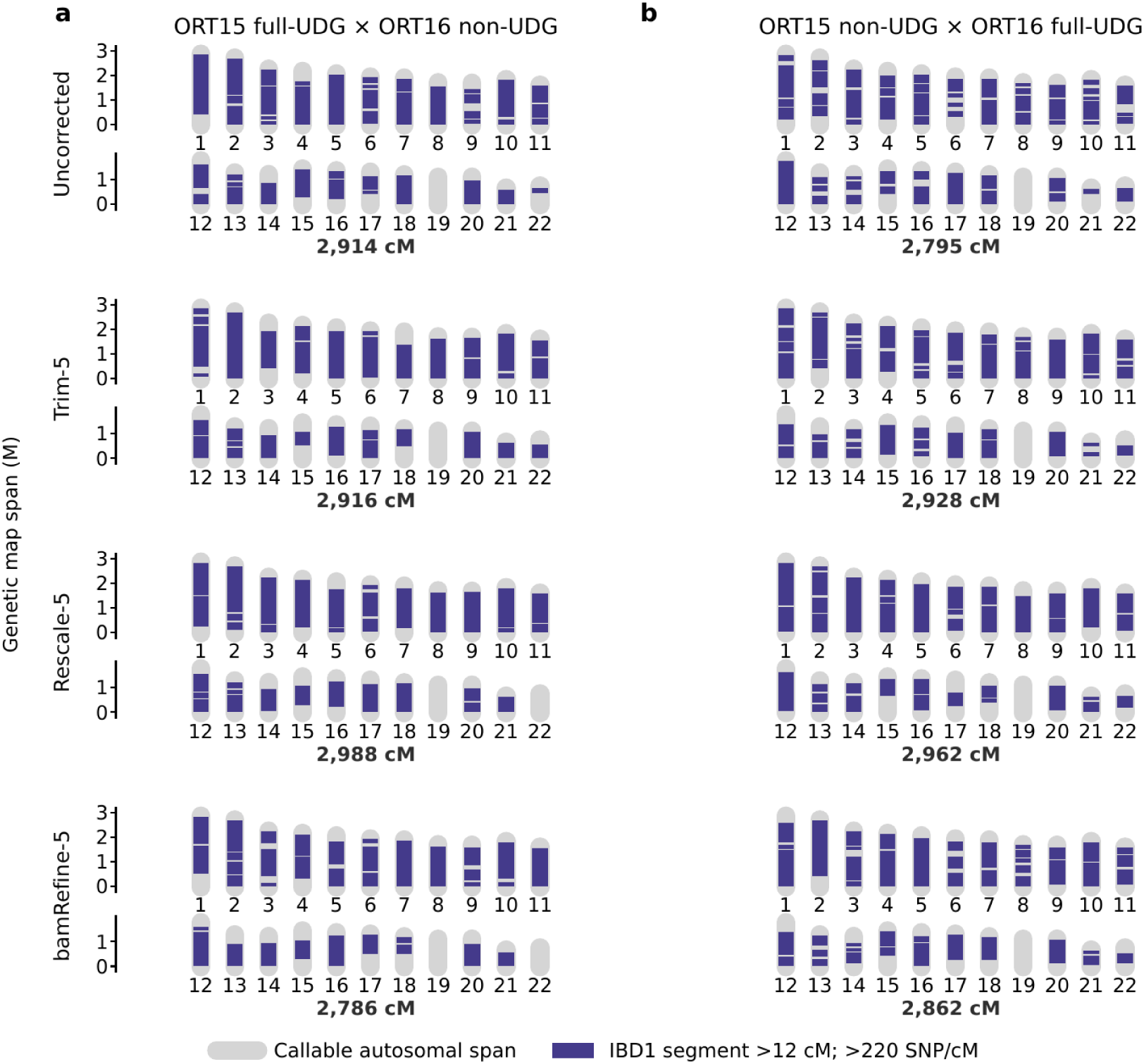
IBD1 karyograms for same-method mixed-library comparisons. Panel **(a)** pairs ORT15 full-UDG with ORT16 non-UDG; panel **(b)** reverses the library types. Within each row, the same correction state is applied to both libraries. Grey bars show the callable autosomal span, and purple intervals show IBD1 segments longer than 12 cM with marker density above 220 SNPs/cM. The bold value below each row is the summed IBD1 length, rounded to the nearest cM. Grey gaps mean that no segment passed these filters; they do not establish absence of biological sharing. Because transition SNPs are included and independent IBD truth is unavailable, the figure assesses whether IBD1 sharing persists across states rather than ranking correction accuracy.

Overall, extensive IBD1 sharing persisted across processing states. However, differences in summed IBD1 length do not identify the most accurate correction method because the analysis included damage-prone transitions and lacked an independent truth set for IBD segments.

### Kinship analyses supported first-degree relatedness, with READv2 and IBS0 favouring a parent-offspring relationship

TKGWV2 reports a half-relatedness coefficient (HRC), with values above 0.1875 classified as first-degree and values from 0.0625 to 0.1875 classified as second-degree. All four full-UDG × full-UDG comparisons were classified as first-degree (HRC, 0.210–0.244; Supplementary Table S6). In both mixed-library directions, the uncorrected comparisons were classified as second-degree (HRC, 0.168 and 0.177), whereas Trim-5, Rescale-5 and bamRefine-5 all produced first-degree classifications (0.201–0.220). When both individuals were represented by non-UDG data, the uncorrected estimate was second-degree (0.142), and the corrected estimates clustered near the first-/second-degree boundary: Trim-5 and bamRefine-5 remained just below it (0.177 and 0.185), whereas Rescale-5 was just above it (0.193). Thus, uncorrected non-UDG data shifted TKGWV2 toward more distant relatedness. Correction restored first-degree classification in the mixed-library comparisons, but classification remained sensitive to processing method when both inputs were non-UDG. Figure 6a shows the full-UDG × full-UDG bamRefine-5 result (HRC = 0.236) as an illustrative TKGWV2 degree estimate, not as a ranking of processing methods; the complete 16-state series is reported in Supplementary Table S6.

**Figure 6.**
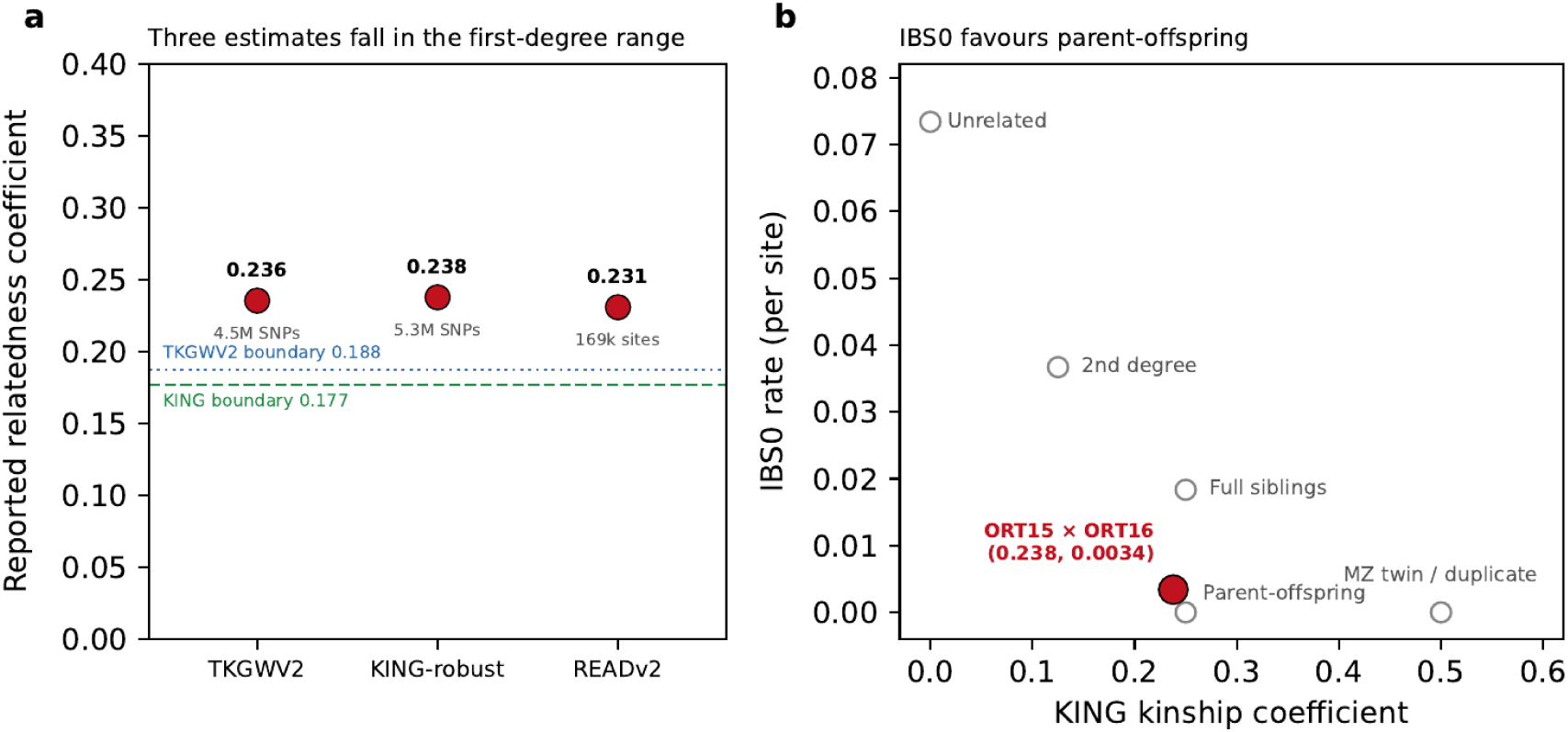
Kinship of ORT15 and ORT16. **(a)** One estimate from each of TKGWV2, KING-robust and READv2 is shown with the first-degree boundaries for TKGWV2 and KING. The TKGWV2 point is the full-UDG × full-UDG bamRefine-5 comparison. The READv2 estimate came from a separate cemetery-wide 1240k call set that included autosomal and sex-chromosome targets; an autosome-only rerun gave the same degree and subtype classification. Because the three estimates use different inputs, they provide complementary degree checks rather than a matched comparison of methods. **(b)** KING-robust kinship and the rate of sites at which the two individuals are opposite homozygotes (IBS0) place the observed pair nearer the parent-offspring expectation than the full-sibling expectation. Together, the estimates support first-degree relatedness, while READv2 and IBS0 favour parent-offspring. Exact values, site counts and sensitivity analyses are reported in Results and Supplementary Table S6.

The primary 1240k panel READv2 analysis classified ORT15 and ORT16 as first-degree with a parent-offspring subtype (KC = 0.231; 169,172 overlapping SNPs). A separate autosome-only 1240k analysis gave the same degree and subtype (KC = 0.232; 157,451 SNPs; Supplementary Table S6). The KING-robust estimate from the uncorrected full-UDG imputed genotypes was 0.238, close to the expected value of 0.25 for first-degree relatives (Fig. 6a). Because KING-robust alone cannot distinguish parent-offspring from full siblings, the subtype interpretation was based on the two READv2 results and the phasing-independent IBS0 comparison.

IBS0 is the fraction of sites at which two individuals are homozygous for opposite alleles. Among 5,272,558 common-variant sites, 17,988 were IBS0, giving a rate of 0.00341. Based on the reference-panel allele frequencies at these sites, the expected unrelated rate was 0.07341. Under the standard outbred Hardy–Weinberg model, full siblings are IBD0 with probability one quarter at an autosomal locus, and IBS0 can occur only in this state.[15] The corresponding theoretical full-sibling expectation was therefore 0.01835. The observed rate was 0.19 times this expectation (Fig. 6b).

As an exploratory sensitivity analysis on chromosome 1, we required a posterior genotype probability of at least 0.99 in both individuals. This reduced IBS0 to 36 of 324,285 sites, or 0.00011. A separate same-individual control comparing nested ORT15 full-UDG read sets yielded 5 IBS0 sites among 408,169 filtered sites, or 0.0000123. Because these analyses used different eligible-site sets, their rates are not directly comparable. The chromosome-1 analyses were supporting checks rather than the primary basis for subtype classification. The genome-wide IBS0 result, together with the READv2 results, favoured parent-offspring over full-sibling relatedness.

### Mitochondrial evidence favoured a mother-son relationship

Both individuals were assigned to mitochondrial haplogroup D4o1, with HaploGrep 3 quality scores of 0.891 for ORT15 and 0.911 for ORT16. ORT15 had 36 called mtDNA SNPs and ORT16 had 37; all 36 ORT15 calls matched ORT16 at the same position and allele (Supplementary Table S7). The sole additional ORT16 call was supported by one read and was not treated as a confirmed difference. The concordant call sets were consistent with shared maternal ancestry. X-to-autosome coverage ratios classified ORT15 as male and ORT16 as female (0.508 and 1.007, respectively; see Methods). Given their parent-offspring relationship, the mitochondrial evidence favoured ORT16 as the mother and ORT15 as the son, although a father-daughter relationship could not be excluded.

### Principal component analysis placement was similar across the tested processing states

We next examined whether PMD correction altered broad population placement by comparing variation across processing states (uncorrected, trimmed, rescaled or masked) and between genotype representations (imputed or pseudo-haploid; Fig. 7). All projections fell within the same broad region of PC1–PC2, near Yakut and other Inner Asian and Siberian reference populations. We measured spread across processing states as the largest pairwise Euclidean distance in PC1–PC2 within each individual, library type and genotype representation. Across four groups, comprising the full-UDG and non-UDG data for each individual, the median spread was 0.00047 for imputed genotypes and 0.00217 for pseudo-haploid calls. The largest spreads were 0.00102 and 0.00336, respectively.

**Figure 7.**
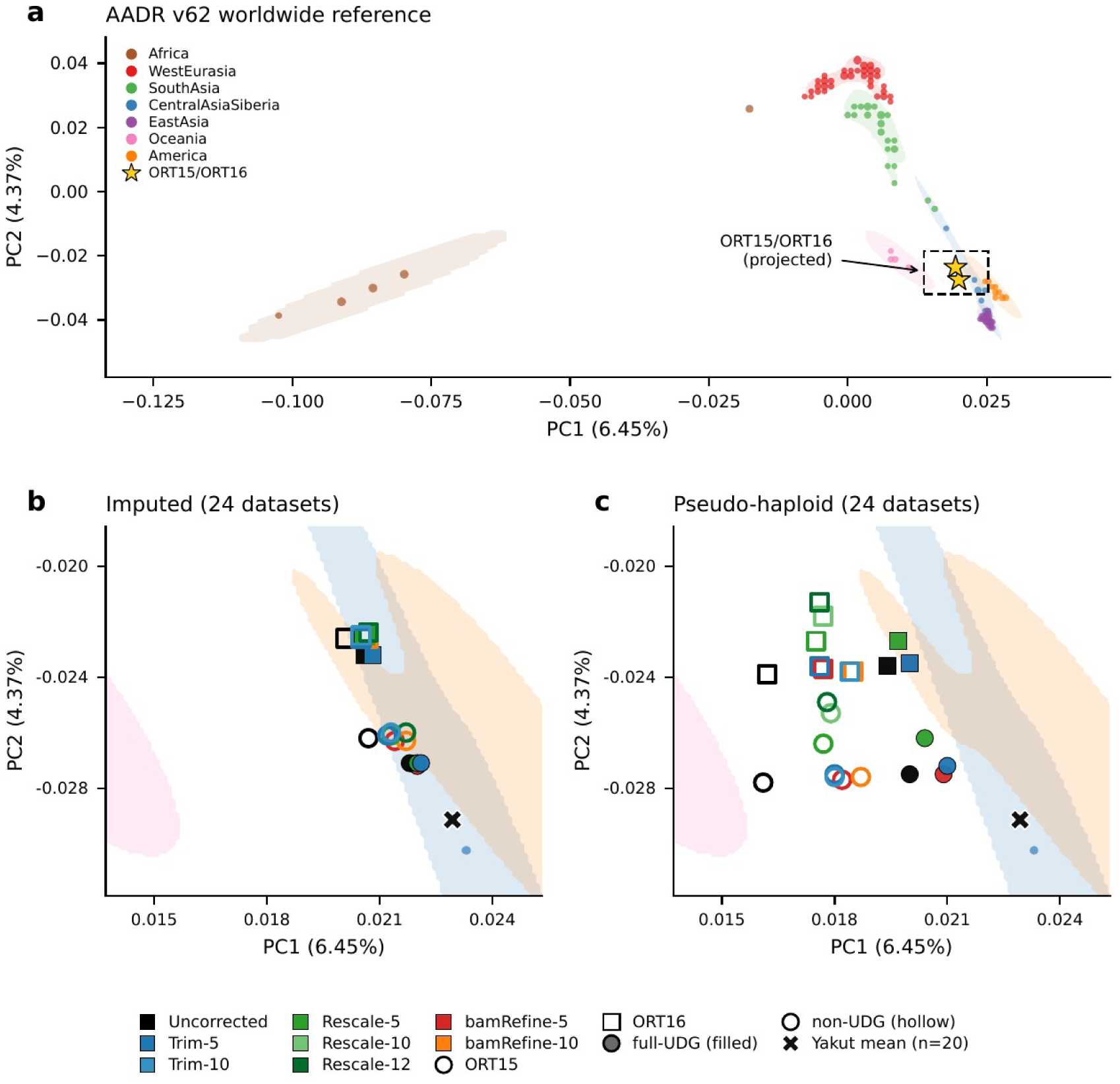
Broad PCA position across PMD correction states. Twenty-four primary datasets were each represented as GLIMPSE-imputed diploid genotypes and non-imputed pseudo-haploid calls and projected onto the same present-day HGDP/SGDP reference axes. The AADR v62.0 reference comprised 985 present-day HGDP/SGDP individuals from 154 populations. The aligned reference-query intersection contained 1,134,702 sites; after internal filtering, 1,113,348 autosomal markers entered smartpca. The set comprised 12 datasets per individual: four full-UDG states and eight primary non-UDG states. The additional 3′-only-5 and 3′-only-10 sensitivity states were not included. **(a)** Worldwide view; gold stars mark representative uncorrected full-UDG pseudo-haploid projections for ORT15 and ORT16 near Yakut in the Northeast Asian and Siberian reference region. **(b,c)** Enlarged views of the imputed and pseudo-haploid datasets on identical axes. Reference samples are coloured by continent; filled density contours are descriptive, not confidence regions, and the Yakut mean is marked. Imputed projections clustered roughly four times more tightly than pseudo-haploid projections. At these coverages, PCA provides a low-resolution check of broad position and does not distinguish among correction methods. Axis percentages use the full eigenvalue trace (PC1 6.45%, PC2 4.37%); eigenvector signs are arbitrary.

By comparison, the imputed and pseudo-haploid projections of the same dataset were separated by a median distance of 0.00309. This distance was 6.6 times the median processing-state spread for imputed genotypes and 1.4 times that for pseudo-haploid calls. Median distances between representations therefore exceeded median spreads across processing states. These dimensionless PCA distances were descriptive; no formal displacement test was performed.

In a separate pseudo-haploid coverage titration, the point clouds generally contracted as coverage increased and were usually tighter for full-UDG than non-UDG datasets. For the comparison between corrected and uncorrected data, seven correction settings were evaluated at seven target coverages in both individuals, giving 98 planned combinations. ORT15 non-UDG Trim-10 at 0.20× could not be generated because its source coverage was 0.173×. For each of the remaining 97 combinations, we compared the mean PCA position of the corrected non-UDG replicates with that of the uncorrected replicates at the same target coverage, using the individual’s uncorrected full-UDG projection at its original coverage as the reference. The corrected mean position was closer to this reference in 87 of the 97 combinations. This titration was descriptive and did not establish ancestry accuracy or identify a coverage threshold (Supplementary Figs. S3 and S4).

## Discussion

These within-individual comparisons show that PMD correction involves trade-offs rather than a single best method. The methods had different effects on genotype agreement and data retention. NRD measures discordance between imputed genotypes obtained from the two library types; lower NRD indicates closer agreement. Compared with the uncorrected non-UDG data, Trim-5 and Trim-10 increased NRD in ORT15 and ORT16, whereas every tested rescaling and masking setting reduced it in both individuals. Rescale-5 and bamRefine-5 produced nearly identical NRD values. In the paired jackknife analysis, which omitted one autosome at a time, the 95% confidence intervals for their NRD difference included zero for ORT15 and ORT16.

Thus, no NRD difference was detected, although the analysis did not establish that the methods were equivalent. Under the conditions tested, rescaling and masking most consistently increased cross-library genotype agreement.

Under this asymmetric damage profile, removal of the relatively low-damage 5′ sequence contributed materially to information loss. Damage in the ORT non-UDG libraries was strongly asymmetric: terminal 5′ C→T was 2.60–3.59%, whereas terminal 3′ G→A was 23.92–26.02%. Because trimBam removed equal lengths from both ends in the primary analysis, it also discarded relatively low-damage 5′ sequence. In the window-matched analysis, 3′-only-5 clipped the same five-base 3′ window as symmetric Trim-5 but left the 5′ end intact. It retained 8.72 and 8.55 percentage points more covered SNPs and yielded observed NRD values 0.82 and 0.67 percentage points lower in ORT15 and ORT16, respectively, while leaving higher ΔALT in both individuals, with confidence intervals excluding zero. Because 3′-only-5 also clipped fewer bases overall, the equal-total-bases comparison provides the more direct test of how clipping was distributed between read ends. Relative to symmetric Trim-5, 3′-only-10 retained 2.16 and 2.44 percentage points more covered SNPs and yielded NRD values 0.60 and 0.40 percentage points lower. Its paired ΔALT differences were small and their confidence intervals included zero, although equivalence was not established.

NRD, covered-site retention, and ΔALT captured non-redundant consequences of processing. Trim-5 and Trim-10 increased NRD while reducing the number of covered SNPs. By contrast, Rescale-5 and bamRefine-5 produced similar NRD estimates despite retaining 99.3% and 92.2– 92.4% of covered sites, respectively. Because NRD was calculated over method-specific evaluable site sets, this comparison is descriptive rather than site matched. ΔALT provided an additional contrast. A negative ΔALT after rescaling does not by itself show that genuine variants were removed, because the estimate also depends on the SNPs retained in the calculation. Correction methods therefore cannot be ranked from NRD alone: cross-library agreement must be considered alongside retained information and residual damage.

For low-coverage libraries with similarly strong 3′-biased damage, symmetric clipping should not be assumed to be optimal. When the same total of ten bases per read was concentrated at the 3′ end, 3′-only-10 retained more covered SNPs and yielded lower observed NRD than symmetric Trim-5, with no ΔALT difference detected in either individual. Damage-informed end-specific clipping may therefore preserve more usable information than equal clipping from both ends when trimming is required. This balance may differ at higher coverage or with more symmetric damage profiles. The end-specific states were not evaluated in the PCA, IBD, or kinship analyses, so this recommendation is limited to covered-site retention and cross-library imputation agreement in these two libraries and does not establish 3′-only trimming as superior to rescaling or bamRefine.

The IBD analyses produced the same broad conclusion across processing states, although detailed segment totals varied. In the mixed-library karyograms, where both datasets had the same processing state, filtered IBD1 segments were observed on 20 or 21 of the 22 autosomes in every state tested. However, total IBD1 length and the ranking of correction methods varied depending on which library types were paired and how they were processed. Rescale-5 yielded the largest observed total in both directions of the two mixed-library analyses. It also produced the largest total in the non-UDG × non-UDG comparison. By contrast, bamRefine-5 produced the largest total in the full-UDG × full-UDG comparison. These findings show that widespread inferred IBD1 sharing persisted across processing states, but differences in total length cannot identify the most accurate correction method. The analysis included transition sites vulnerable to PMD and lacked known true IBD segments for comparison. We also did not assess phasing accuracy or switch errors, which may have affected segment length and fragmentation.

Kinship estimates fell near the 0.1875 first-degree threshold and shifted across correction states. In the non-UDG/non-UDG comparison, TKGWV2 placed Rescale-5 above the threshold, whereas Trim-5 and bamRefine-5 remained just below it. Because all three estimates were close to the cutoff, the different categorical labels should not be interpreted as correspondingly large differences in relatedness. The broader evidence remained consistent with a first-degree parent-offspring relationship: READv2 and the low IBS0 rate supported this interpretation, while ancIBD identified widespread IBD1 sharing. Thus, the threshold-based label varied in one comparison without changing the overall relationship interpretation.

The kinship analyses provided complementary checks rather than independent replications because they used the same individuals and partly overlapped in their input reads, SNP sets or imputed genotypes. The primary and autosome-only 1240k READv2 analyses classified the ORT15 and ORT16 pair as parent-offspring, and the low genome-wide IBS0 rate also favoured parent-offspring over full-sibling relatedness. The KING-robust estimate supported first-degree relatedness but could not distinguish these two subtypes. READv2 estimates also depend on the normalisation baseline. Because the normalisation cohorts were not screened to remove related pairs, their composition may have influenced the exact coefficients.

The mitochondrial evidence favoured a mother-son relationship but did not exclude father-daughter relatedness. Both individuals were assigned to mitochondrial haplogroup D4o1, and no well-supported mtDNA SNP difference was identified between them. Given their parent-offspring relationship and genetic-sex estimates, this evidence favoured ORT16 as the mother and ORT15 as the son. Fathers do not transmit mtDNA. If ORT15 were ORT16’s father, ORT16 would instead have inherited her mtDNA from her mother, who would also have needed to carry a mitochondrial profile closely matching that of ORT15.

PCA provided a low-resolution check of broad population placement. Across the titration analysis, 87 of 97 comparisons shifted toward the matched full-UDG position. At this coverage, however, PCA has limited resolution for smaller changes; therefore, an unchanged broad placement cannot establish that processing had no effect. The analysis describes the direction of the observed coordinate shifts but does not distinguish among correction methods.

The matched same-extract design reduced variation attributable to individual and source material, but library type, preparation, and sequencing depth remained intertwined. For each individual, the full-UDG comparator pooled three library reactions and had 3.3–4.2-fold greater coverage than the single non-UDG library. This greater depth could increase genotype certainty and the number of evaluable loci in the full-UDG data. The respective contributions of treatment, preparation, and depth to absolute cross-library differences therefore cannot be separated here. Comparisons among correction states derived from the same non-UDG starting BAM are not affected by this between-library depth confound, although usable depth and eligible-site composition change by design. Future studies should generate replicate full-UDG and non-UDG libraries from the same extracts and sequence them to matched depths.

The scope of inference is also limited by the study design. Only two individuals and one parent-offspring relationship were examined, and such close relationships are easier to detect than more distant ones. This is therefore a paired case study rather than a population-scale benchmark. Unrelated and more distant pairs were not imputed and processed through the same PMD-correction and ancIBD design. The non-UDG coverages were near the lower range reported for ancIBD,[14] and the inferred totals remained conditional on the marker intersection, segment filters and untested phasing accuracy. Coverage was titrated for PCA but not for imputation or IBD. Furthermore, the asymmetric damage pattern associated with this library-preparation workflow may not represent conventional untreated non-UDG libraries. These features limit generalisation to other damage patterns, coverages and relationship classes.

Finally, the 1000 Genomes Phase 3 panel contains no Central Asian, Mongolic or Siberian reference population,[12] limiting fine-scale imputation for the ORT individuals. Their broad PCA placement was nevertheless also observed in the non-imputed pseudo-haploid data. Future studies should test more representative resources, including the harmonised HGDP plus 1000 Genomes panel[16] and a Northeast Asian panel such as NARD.[17] They should also include conventional non-UDG libraries, matched-depth comparisons, larger sample sets and several relationship classes. Testing end-specific and damage-adaptive trimming across additional libraries spanning broader coverage and damage profiles would help determine whether the improvements in covered-site retention and cross-library concordance observed here generalise beyond these strongly asymmetric, low-coverage libraries.

## Conclusions

Computational PMD correction changed damage-associated allele support, covered-site retention, imputation agreement and inferred IBD1 length in these two low-coverage ancient genomes. bamRefine-5 and Rescale-5 differed by only 0.02–0.03 percentage points in NRD; the paired chromosome-jackknife confidence intervals spanned zero, providing no evidence of a difference but not establishing equivalence. In non-UDG data, Rescale-5 retained 99.3% of covered sites, whereas bamRefine-5 retained about 92% and left a small positive ΔALT. IBD1 remained widespread across the autosomes. READv2 and KING supported first-degree relatedness, while TKGWV2 classifications depended on library type and correction state. READv2 and IBS0 favoured a parent-offspring relationship. Maternal-lineage evidence favours ORT16 as the mother and ORT15 as the son, although father-daughter cannot be excluded. Because the study includes one pair and no independent genotype truth, it demonstrates method trade-offs rather than a universal ranking.

## Methods

### Archaeological material and radiocarbon dating

ORT15 and ORT16 came from separate graves at the Oortsog Ovoo burial ground in Tuvshinshiree soum, Sukhbaatar Province, eastern Mongolia (Supplementary Figure S5). The 2021 excavation report dates the cemetery broadly to the 13th and 14th centuries CE.[18] ORT15 was Burial 4 in the centre of the cemetery, and ORT16 was Burial 8 toward its northeastern edge. Both were stone-covered graves that were intact or largely intact at excavation. The field report described Burial 4 as male and Burial 8 as female. Administrative boundaries in Supplementary Figure S5 were obtained from geoBoundaries.[19]

After filtering reads to mapping quality ≥30, X-to-autosome read-density ratios averaged across the 22 autosomes identified ORT15 as genetically male (0.508) and ORT16 as genetically female (1.007).[20] The pair was selected because preliminary analyses indicated first-degree relatedness, allowing the same relationship to be examined across matched non-UDG and full-UDG libraries after different PMD corrections. The remains are housed at the Chinggis Khaan National Museum, Ulaanbaatar. Pre-sampling micro-CT images of the petrous bones are shown in Supplementary Figure S6.

For radiocarbon dating, 212.2 mg of ORT16 petrous bone was used for collagen extraction, and the collagen was dated at Paleo Labo Co., Ltd. The measured ¹⁴C age was 705 ± 15 yr BP. The laboratory reported a principal calibrated 2σ range from 1274 to 1299 cal AD (92.62%) and a minor range from 1372 to 1376 cal AD (2.83%). ORT16 therefore most probably dates to the late 13th century CE (Supplementary Table S8). These calendar ranges were reproduced from the laboratory age table because the calibration software and curve were not recorded in the materials available for this study. ORT15 was not radiocarbon dated.

### Ancient DNA extraction and sequencing

Sampling, DNA extraction and library preparation were performed in a dedicated ancient-DNA clean room.[21] Before sampling, micro-CT was used to locate the cochlea (Supplementary Figure S6). After 30 min of UV irradiation (254 nm, 7000 mJ/cm^2^), we collected 26.5 mg of cochlear bone powder from ORT15 and 36.2 mg from ORT16.

To reduce surface contamination, the powder was pre-digested for 15 min at 37°C in 1 mL lysis buffer without Proteinase K (Thermo Scientific, EO0492). The supernatant was retained at −80°C, and the pellet was then digested for 16 h at 37°C in 1 mL lysis buffer with Proteinase K. DNA was extracted from 150 µL lysate with an automated silica-coated magnetic-bead method[22] (G-Biosciences, 786-917). We reduced 2-propanol in binding buffer D from 40% to 20% (vol/vol) to limit retention of very short fragments. Extracts were eluted in 40 µL per well, and wells from the same individual were pooled for library preparation.

Double-stranded libraries were prepared with the NEBNext Ultra II DNA Library Prep Kit (New England Biolabs, E7645L). For each individual, four aliquots of 15 µL from the same pooled extract were used for three separately indexed full-UDG reactions (a–c) and one non-UDG reaction. For the full-UDG libraries, DNA extracts underwent dedicated USER treatment using the full-UDG incubation protocol before library preparation. This treatment was performed for 3 h at 37°C. The non-UDG reaction omitted this pre-library treatment. After adaptor ligation, however, all libraries were incubated with USER enzyme (New England Biolabs, M5505L) for 15 min at 37°C to open the uracil-containing NEBNext loop adaptor. T4 polynucleotide kinase (M0201S) and T4 DNA polymerase (M0203S) were also used for the full-UDG reactions, followed by MinElute purification (Qiagen, 28004). The three full-UDG reactions were merged downstream.

All libraries underwent 12 indexing-PCR cycles with NEBNext Multiplex Oligos for Illumina (Sets 2–4; New England Biolabs, E6442, E6444 and E6446). Libraries were cleaned with SPRIselect beads (Beckman Coulter, B23318), quantified with the Qubit 1× dsDNA HS assay (Invitrogen, Q33231), and checked with the Agilent High Sensitivity D1000 TapeStation assay (5067-5585). After screening on an Illumina iSeq 100 for endogenous human DNA and terminal damage, the libraries were sequenced by Novogene on an Illumina NovaSeq X with 2×150-bp reads. An extraction blank was carried through extraction, library preparation and iSeq 100 screening.

### DNA sequence data processing

Adapters were removed, terminal bases at or below quality 25 were trimmed, and overlapping pairs were collapsed with AdapterRemoval. Reads shorter than 30 bp after trimming were discarded.[23] We compared three read sets: collapsed reads alone; collapsed plus truncated-collapsed reads; and those reads plus unmerged pairs. Reads were mapped to hs37d5 with BWA aln; BAMs were merged across sequencing runs and deduplicated with samtools.[24,25] Endogenous human DNA proportion and mean genomic coverage were summarised with Qualimap.[26] All three read sets were evaluated for coverage and endogenous content, but the imputation and IBD analyses used collapsed reads alone because the additional categories provided little coverage gain and unmerged pairs lowered the endogenous DNA proportion.

Detailed preprocessing and duplicate-removal commands are provided in the Supplementary Code. Software and external-resource versions are listed in Supplementary Table S9.

### Contamination estimation

We estimated present-day human DNA contamination using two complementary approaches (Supplementary Table S2). Nuclear contamination was assessed from apparent X-chromosome heterozygosity in genetically male ORT15. Because a male is expected to be hemizygous across most of chromosome X, excess support for a second allele can indicate contamination after sequencing error is modelled. This approach requires X-chromosome hemizygosity and therefore does not provide an equivalent estimate for genetically female ORT16. We used ANGSD on the collapsed full-UDG BAM across 6757 covered HapMapChrX sites and estimated contamination from apparent X-chromosome heterozygosity by method of moments and maximum likelihood.[27,28]

Mitochondrial contamination was assessed in both individuals with mtCont (schmutzi), which compares the observed mitochondrial reads from the collapsed, deduplicated non-UDG BAM against the Eurasian haplotype panel.[29] Because these libraries carry asymmetric damage (reduced 5′ C→T, strong 3′ G→A, consistent with USER exposure during library preparation), we supplied separately estimated 5′ and 3′ damage profiles rather than a symmetric one, with endogenous-base likelihoods from endoCaller. The analysis assumed a single contaminant haplotype and was run standalone, not as the full iterative schmutzi workflow. Commands, filters and resource files are given in the Supplementary Code; software versions are listed in Supplementary Table S9.

### PMD correction strategies

For each library, we retained the standard-processed BAM without computational PMD correction (hereafter “uncorrected”) and generated corrected BAMs with three approaches. For the damage-signal and covered-site retention analyses, the full-UDG series comprised uncorrected, Trim-5, Rescale-5, Rescale-12 and bamRefine-5. The primary non-UDG series comprised eight states: uncorrected, Trim-5, Trim-10, Rescale-5, Rescale-10, Rescale-12, bamRefine-5 and bamRefine-10. trimBam soft-clipped the stated number of bases from each read end; these bases remain in the BAM but do not contribute to genotype likelihoods. mapDamage2 lowered the quality of bases likely to be damaged, using the stated terminal modelling length; Rescale-12 used its default 12-bp setting. bamRefine masked likely damaged bases that overlapped biallelic 1000 Genomes SNPs. Damage profiles were checked with mapDamage2 after correction (Supplementary Figure S2); recorded parameters and portable workflow scripts are provided in the Supplementary Code and Data.

As an additional sensitivity analysis, we generated 3′-only-5 and 3′-only-10 states from each collapsed, deduplicated non-UDG BAM by clipping only the strand-aware 3′ end and leaving the 5′ end unchanged. This clipped the right side of forward-strand alignments and the left side of reverse-strand alignments. The input BAMs contained only collapsed reads and no alignments marked as paired. We compared 3′-only-5 with symmetric Trim-5 over the same five-base 3′ window, and 3′-only-10 with symmetric Trim-5 while holding the total number of clipped bases per read at ten. All four derived BAMs retained the input alignment counts and were processed through the same pileup, ΔALT, imputation and concordance workflows as the primary non-UDG states, using the same uncorrected full-UDG comparator for each individual. Commands, software versions and integrity checks are provided in Supplementary Table S9 and the Supplementary Code.

### Damage-associated signal and covered-site retention

To assess damage-associated allele support, we examined biallelic SNPs from the 1000 Genomes Project.[12,25] At sites with combined reference and alternate read depth (REF+ALT) ≥3, we calculated the site-level alternative-allele fraction. We calculated the mean separately for damage-prone C→T/G→A changes and transversions. At each eligible site, the alternative-allele fraction p was ALT/(REF+ALT). We defined ΔALT as follows:

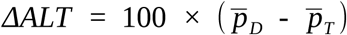

Here, D denotes damage-prone C→T and G→A SNPs, T denotes transversions, and the bars denote the mean site-level alternative-allele fraction within each class. Multiplication by 100 expresses ΔALT in percentage points. Positive values indicate greater alternative-allele support at damage-prone sites. Because the SNP classes can differ in allele frequency and reference or alternative allele composition, and because correction states can retain different sites, ΔALT is descriptive rather than a direct estimate of PMD or genuine variant loss. We estimated uncertainty in ΔALT and the prespecified differences between correction states with jackknives over genomic regions. The primary analysis omitted one autosome at a time; sensitivity analyses used fixed 5-Mb and 10-Mb blocks across the GRCh37 autosomes. In each replicate, the same region was omitted from every state in the paired comparison, and ΔALT was recalculated from the class-specific sums of site-level alternative-allele fractions and eligible-site counts. Jackknife standard errors and bias-corrected, two-sided 95% confidence intervals used a t distribution with B − 1 degrees of freedom, where B is the number of blocks. A further sensitivity analysis included only sites whose chromosome, position, reference allele and alternative allele matched across every state in the comparison and whose combined REF+ALT depth was ≥3 in every state. These intervals describe variation among genomic regions in the analysed libraries; they do not estimate variation among individuals.

Covered-site retention was calculated as the percentage of SNPs with combined REF+ALT depth ≥1 after correction, relative to the uncorrected library from the same individual and library type. It measures how many covered SNPs remained under the stated mapping and base quality filters; it does not measure read depth at those sites or genotype accuracy. Exact filters and commands are provided in the Supplementary Code.

### Genotype imputation and cross-library agreement

For autosomal SNPs, we calculated genotype likelihoods at biallelic sites in the 1000 Genomes Phase 3 panel with bcftools. We required mapping quality ≥30 and base quality ≥20, excluded indels and did not require a minimum read depth.[12,25] This differs from the preceding allele-fraction analysis, which required a depth of three reads. Each dataset was then imputed and phased independently with GLIMPSE using the Phase 3 SHAPEIT2 integrated v5b reference panel of 2504 samples and GRCh37 genetic maps.[13] We did not evaluate phasing accuracy or switch errors; the phased output was used only for downstream IBD analysis.

For each individual, we compared ten non-UDG datasets with one fixed comparator: the independently imputed full-UDG dataset without computational damage correction. The ten non-UDG states were uncorrected, Trim-5, Trim-10, 3′-only-5, 3′-only-10, Rescale-5, Rescale-10, Rescale-12, bamRefine-5 and bamRefine-10. Because this comparator is another library from the same individual, agreement does not show whether either genotype is correct. We assessed agreement with the GLIMPSE2 concordance module using dosage *r*², best-guess *r*², and non-reference discordance (NRD).[30] Dosage *r*² is based on the expected alternate-allele dosage from genotype probabilities; best-guess *r*² is based on the most probable genotype. The two *r*² summaries were weighted by the number of variants in each minor-allele-frequency (MAF) bin and restricted to MAF ≥0.1%. NRD included all frequency bins and was the fraction of evaluable comparisons involving a non-reference genotype in which the non-UDG dataset and full-UDG comparator disagreed.

The NRD difference was defined as bamRefine-5 minus Rescale-5. Negative values indicate lower NRD for bamRefine-5; positive values indicate lower NRD for Rescale-5. We estimated uncertainty with a paired jackknife that omitted one autosome at a time. In each of 22 replicates, the same autosome was removed from both methods before the difference was recalculated.

Because the two methods could retain different sites, pairing was by omitted autosome rather than by individual site. We report the observed difference, bias-corrected jackknife estimate, jackknife standard error and 95% confidence interval from a t distribution with 21 degrees of freedom. Full imputation, concordance and jackknife procedures are provided in the Supplementary Code; software versions are listed in Supplementary Table S9.

### IBD inference

Phased GLIMPSE VCFs were analysed with ancIBD, which uses a hidden Markov model to follow imputed haplotypes along the genome and identify regions inherited from a recent common ancestor.[14] The non-UDG libraries (0.27× and 0.35×) were close to the program’s approximate 0.25× lower coverage range for whole-genome data, so calls were interpreted cautiously. We report IBD1: regions where the pair is inferred to share one chromosome copy by descent. IBD1 can occur in several relationship classes and is therefore not, by itself, specific to first-degree relatives.

The analysis used 1,100,313 autosomal SNPs whose chromosome, position, reference allele and alternate allele matched across the Allen Ancient DNA Resource (AADR) v62.0 targets, 1000 Genomes reference-frequency records and the imputed data.[31,32] AADR v62.0 supplied genetic positions in Morgans. ancIBD initially called segments from 8 centimorgans (cM; 0.08 Morgan), but segments shorter than 12 cM are more vulnerable to false positives in low-coverage data. Our primary summary therefore retained segments strictly longer than 12 cM and denser than 220 SNPs/cM; the density filter removes calls in sparsely marked regions.[14] We also summarised segments longer than 8, 16, and 20 cM. Segments were not externally gap-merged, and IBD2 was not used. We recorded retained-segment count and summed IBD1 length relative to the callable autosomal span of 3538.85 cM, defined as the summed first-to-last marker distance across the autosomes.

We used two focused comparison designs, each based on four states: uncorrected, Trim-5, Rescale-5 and bamRefine-5. In the *asymmetric* design, one individual’s full-UDG library remained uncorrected while only the other individual’s non-UDG state varied, in both directions. This design isolates changes on one side of the pair. In the *same-method* design, the same state was applied to both libraries across the full-UDG × full-UDG, non-UDG × non-UDG, and two directional mixed-library pairings. The mixed directions were kept separate because the individual, library type and coverage differed on each side. The complete 8×8 matrix additionally contains every cross-state pairing; those extra cells are diagnostic and were not used to attribute an effect to one method. The complete matrix is shown in Supplementary Figure S7, and the relationship between total segment length and segment count is shown in Supplementary Figure S8. Implementation details are provided in the Supplementary Code.

### Kinship estimation

Relationship degree and subtype were evaluated with four complementary summaries: TKGWV2, READv2, KING-robust kinship and the rate of sites at which the pair is opposite homozygous (IBS0). They are not fully independent because some share reads, SNP ascertainment or imputed genotypes; KING and IBS0 use the same merged diploid calls.

TKGWV2 used pseudo-haploid genotypes from the collapsed BAMs and 1000 Genomes Phase 3 East Asian allele frequencies.[33] Its half-relatedness coefficient (HRC) classified values below 0.0625 as unrelated, 0.0625–0.1875 as second-degree, and values above 0.1875 as first-degree. We examined full-UDG, non-UDG and directional mixed-library pairs in the uncorrected, Trim-5, Rescale-5 and bamRefine-5 states (Supplementary Table S6).

The primary READv2 analysis evaluated ORT15 and ORT16 within an archived cemetery-wide 1240k call set. For ORT15 and ORT16, the input BAMs combined full-UDG libraries a–c and were soft-clipped by 6 bp at each read end. Pseudo-haploid calls included autosomal and sex-chromosome targets. Twenty-one retained individuals generated 210 pairwise comparisons, whose median pairwise mismatch proportion (P_0_) provided the primary normalisation baseline. An autosome-only 1240k rerun used an archived call set containing 22 individuals.[8] READv2 reports a normalised kinship coefficient (KC) and a first-degree subtype. Only the result for the ORT15 and ORT16 pair is interpreted here. The public reproducibility repository provides the anonymised 210-pair P_0_ table, focal outputs, downstream READv2 commands, the normalisation procedure and validation records. Full cemetery genotype matrices and the controlled BAM inputs are not publicly distributed. The additional individuals were not included in the PMD-correction, imputation, IBD or PCA analyses. The normalisation cohorts were not filtered to exclude related pairs, and no relative-excluded analysis was performed; estimates may therefore depend on cohort composition. The exact original random state used during the primary pseudo-haploid sampling step was not preserved, so a new BAM-level run need not be byte-identical to the archived analysis.

We calculated KING-robust kinship and IBS0 from the same merged, uncorrected full-UDG diploid genotypes imputed with GLIMPSE.[15] A KING value near 0.25 indicates first-degree relatedness. IBS0 counts sites where the two individuals are opposite homozygotes; such sites should be nearly absent in parent-offspring pairs but remain present in full siblings. The genome-wide analysis used 5,272,558 variants with reference-panel allele frequencies from 0.1 to 0.9. To assess residual imputation error, we repeated the analysis on chromosome 1 using genotypes with posterior probability ≥0.99 in both individuals (324,285 sites). Formulas and implementation details are provided in the Supplementary Code.

### Mitochondrial haplogroup assignment

For each individual, mitochondrial variants were called from deduplicated full-UDG data containing collapsed reads from libraries a–c. The alternative overlapping read sets were not used. Reads were aligned with BWA to hs37d5, which includes nuclear, decoy and rCRS mitochondrial sequences (NC_012920).[24,34] Mitochondrial variants were called as haploid with bcftools. We required mapping and base qualities of at least 30, retained the default maximum depth of 250 reads per file and did not impose global minimum-depth, allele-fraction or strand-balance thresholds.[25] SNPs were compared only when the chromosome, position and both alleles matched; a missing call was treated as missing rather than as a reference genotype. The MT:3106 CN>C record was excluded because it represents the artificial rCRS spacer at position 3107 rather than biological variation. Haplogroups were assigned with HaploGrep 3 using the PhyloTree 17 Forensic Update.[35–37] Scores were interpreted alongside the underlying variant calls, without applying a universal cutoff. Software versions, commands and full processing details are provided in Supplementary Table S9 and the Supplementary Code.

### Population genetic context

To test whether PMD correction changed broad population placement, we projected 24 datasets, 12 per individual, in two forms. The four full-UDG states were uncorrected, Trim-5, Rescale-5 and bamRefine-5. The eight primary non-UDG states comprised the same four plus Trim-10, Rescale-10, Rescale-12 and bamRefine-10. One form used GLIMPSE-imputed genotypes; the other used non-imputed pseudo-haploid calls made by sampling one observed allele at each covered 1240k site. This produced 48 projected points on the same reference axes.

The axes were built with smartpca from EIGENSOFT build 18140, using 985 present-day individuals in the Human Genome Diversity Project (HGDP) and Simons Genome Diversity Project (SGDP), representing 154 populations in AADR v62.0; ancient and archaic individuals were excluded.[31,38–41] After allele alignment and internal filtering, 1,113,348 autosomal markers were analysed. ORT datasets were projected without influencing the axes, and no LD pruning or outlier-removal iterations were applied. PC1–PC4 explained 6.45%, 4.37%, 1.53%, and 1.04% of total variance, calculated against all 984 non-zero components. We described correction-associated spread as the largest pairwise PC distance within each individual, library type and genotype representation, and compared it with scatter among 20 Yakut reference individuals.

For the separate coverage titration, the same 24 correction-state BAMs were downsampled to 0.01, 0.02, 0.04, 0.06, 0.08, 0.10 and 0.20× where attainable. Source depth was measured with samtools (Supplementary Table S10), and the retained fraction was target depth divided by source depth. Ten random subsamples were generated per attainable target with replicate seeds 1–10; targets above the source depth were omitted. The BAMs were restricted to 1240k sites before uniform read subsampling, and one observed allele per covered site was sampled with pileupCaller.[42] The resulting pseudo-haploid genotypes were projected in a separate smartpca run without influencing the reference axes. The final dataset contained 1694 projected points; the ten ORT15 non-UDG Trim-10 replicates at 0.20× were absent because that source library had lower coverage. We interpreted the PC1–PC2 point clouds descriptively and did not apply a formal displacement threshold. Exact parameters, marker traces and formulas for both PCA analyses are provided in the Supplementary Code.

### Ethics, permissions and sample curation

The 2021 excavation at Oortsog Ovoo was conducted under the required archaeological authorisation.[18] The remains are curated by the Chinggis Khaan National Museum. Under a Memorandum of Understanding between the museum and Kanazawa University, the museum authorised the temporary transfer of the specimens to Japan for minimally destructive sampling and ancient DNA analysis; the specimens were subsequently returned to the museum. All procedures complied with the agreement, the museum’s requirements and applicable Mongolian and Japanese regulations. No separate project-specific permit or ethics approval number was assigned. Informed consent was not applicable.

### Use of generative AI and AI-assisted technologies

The authors used Claude (Anthropic) and Codex (OpenAI) for language editing and assistance with refining and validating analysis scripts. All AI-assisted revisions were reviewed by the authors, who take full responsibility for the article. No generative image models were used.

## Code availability

Workflow scripts, supporting data and validation materials (Supplementary Code and Data), including anonymised pairwise P_0_ tables and READv2 validation outputs, are available in GitHub release v1.2.2 (https://github.com/Tsenomiu/ORT_PMD_reproducibility/releases/tag/ v1.2.2; commit 888c579) and archived on Zenodo (https://doi.org/10.5281/zenodo.22502035). Author-written code and derived tables use the MIT License; the TKGWV2 patch uses GPL-2.0-only. Software versions are listed in Supplementary Table S9.

## Data Availability

Raw sequencing reads for all eight study libraries and associated runs are publicly available in the European Nucleotide Archive under project accession PRJEB112497. Third-party reference resources, including AADR v62.0, the 1000 Genomes Project panel and hs37d5, are available from their original providers. Genotype and BAM data from additional cemetery individuals used for READv2 normalisation are not publicly deposited because they form part of an unpublished cemetery study. Requests for access should be directed to the corresponding author at.

## Acknowledgements

We gratefully acknowledge the late Dashdulam Dashdeleg for her early support in establishing this project. We thank Solongo Batbold for facilitating access to the museum samples and Dr.Oyunbold Lamid-Ochir at the First Hospital of Mongolia for providing access to CT scanning facilities. All original figures and the workflow schematic were prepared by the authors; the archaeological plan and administrative-boundary sources are credited in Supplementary Figure S5.

## Research funding

This work was supported by JSPS KAKENHI Grant Numbers JP26K18397, JP25H00487, JP21H04358 and JP24H00097, and by the JSPS Program for Forming Japan’s Peak Research Universities (J-PEAKS) Grant Number JPJS00420230006. The SAKIGAKE project at Kanazawa University also supported this work.

## Author contributions

O.R. conceived and designed the study, performed the laboratory work and all computational analyses, prepared the figures, and wrote the original manuscript. C.S. provided access to the skeletal collection and contributed archaeological information and context. K.J. supervised the work conducted in Mongolia and contributed to manuscript review and editing. T.G. provided overall project supervision and scientific guidance, acquired funding, and critically reviewed and edited the manuscript. O.R. also contributed to funding acquisition. All authors reviewed and approved the final manuscript.

## Additional information

### Competing interests

The authors declare no competing interests.

**Correspondence** and requests for materials should be addressed to O.R. or T.G.

**Supplementary information** is available for this paper. It contains Supplementary Figures S1– S8 and Supplementary Tables S1–S10.

## References

[1] Briggs, A. W., et al. Patterns of damage in genomic DNA sequences from a Neandertal. Proc. Natl Acad. Sci. USA 104, 14616–14621 (2007).

[2] Briggs, A. W. et al. Removal of deaminated cytosines and detection of in vivo methylation in ancient DNA. Nucleic Acids Res. 38, e87 (2010).

[3] Meyer, M. et al. A high-coverage genome sequence from an archaic Denisovan individual. Science 338, 222–226 (2012).

[4] Gansauge, M.-T. & Meyer, M. Single-stranded DNA library preparation for the sequencing of ancient or damaged DNA. Nat. Protoc. 8, 737–748 (2013).

[5] Rohland, N., Harney, E., Mallick, S., Nordenfelt, S. & Reich, D. Partial uracil-DNA-glycosylase treatment for screening of ancient DNA. Phil. Trans. R. Soc. B 370, 20130624 (2015).

[6] Koptekin, D. et al. Pre-processing of paleogenomes: mitigating reference bias and postmortem damage in ancient genome data. Genome Biol. 26, 6 (2025).

[7] Garrido Marques, A., Rubinacci, S., Malaspinas, A.-S., Delaneau, O. & Sousa da Mota, B. Assessing the impact of post-mortem damage and contamination on imputation performance in ancient DNA. Sci. Rep. 14, 6227 (2024).

[8] Alaçamlı, E. et al. READv2: advanced and user-friendly detection of biological relatedness in archaeogenomics. Genome Biol. 25, 216 (2024).

[9] Jun, G. et al. An efficient and scalable analysis framework for variant extraction and refinement from population-scale DNA sequence data. Genome Res. 25, 918–925 (2015).

[10] Jónsson, H., Ginolhac, A., Schubert, M., Johnson, P. L. F. & Orlando, L. mapDamage2.0: fast approximate Bayesian estimates of ancient DNA damage parameters. Bioinformatics 29, 1682–1684 (2013).

[11] Hui, R., D’Atanasio, E., Cassidy, L. M., Scheib, C. L. & Kivisild, T. Evaluating genotype imputation pipeline for ultra-low coverage ancient genomes. Sci. Rep. 10, 18542 (2020).

[12] The 1000 Genomes Project Consortium. A global reference for human genetic variation. Nature 526, 68–74 (2015).

[13] Rubinacci, S., Ribeiro, D. M., Hofmeister, R. J. & Delaneau, O. Efficient phasing and imputation of low-coverage sequencing data using large reference panels. Nat. Genet. 53, 120–126 (2021).

[14] Ringbauer, H. et al. Accurate detection of identity-by-descent segments in human ancient DNA. Nat. Genet. 56, 143–151 (2024).

[15] Manichaikul, A. et al. Robust relationship inference in genome-wide association studies. Bioinformatics 26, 2867–2873 (2010).

[16] Koenig, Z. et al. A harmonized public resource of deeply sequenced diverse human genomes. Genome Res. 34, 796–809 (2024).

[17] Yoo, S. K. et al. NARD: whole-genome reference panel of 1779 Northeast Asians improves imputation accuracy of rare and low-frequency variants. Genome Med. 11, 64 (2019).

[18] Erdenebold, L., Erdene, J., Ariunbold, G., Bolormaa, S. & Gantogtokh, G. Report on the 2021 Archaeological Excavation and Research at Ikh Oortsog Mountain, Tuvshinshiree Soum, Sukhbaatar Province (Mongolian University of Science and Technology, Chinggis Khaan Museum and Chinggis Khaan Institute of Heritage and Culture, 2022). Field report, in Mongolian.

[19] Runfola, D. et al. geoBoundaries: a global database of political administrative boundaries. PLoS One 15, e0231866 (2020).

[20] Mittnik, A., Wang, C.-C., Svoboda, J. & Krause, J. A Molecular Approach to the Sexing of the Triple Burial at the Upper Paleolithic Site of Dolní Věstonice. PLoS ONE 11, e0163019 (2016).

[21] Ishiya, K. et al. Towards on-site paleogenomics: application and perspective of nanopore sequencing with ancient DNA. Preprint at 10.64898/2025.12.29.696801 (2025).

[22] Rohland, N., Glocke, I., Aximu-Petri, A. & Meyer, M. Extraction of highly degraded DNA from ancient bones, teeth and sediments for high-throughput sequencing. Nat. Protoc. 13, 2447–2461 (2018).

[23] Schubert, M., Lindgreen, S. & Orlando, L. AdapterRemoval v2: rapid adapter trimming, identification, and read merging. BMC Res. Notes 9, 88 (2016).

[24] Li, H. & Durbin, R. Fast and accurate short read alignment with Burrows-Wheeler transform. Bioinformatics 25, 1754–1760 (2009).

[25] Danecek, P. et al. Twelve years of SAMtools and BCFtools. GigaScience 10, giab008 (2021).

[26] Okonechnikov, K., Conesa, A. & García-Alcalde, F. Qualimap 2: advanced multi-sample quality control for high-throughput sequencing data. Bioinformatics 32, 292–294 (2016).

[27] Korneliussen, T. S., Albrechtsen, A. & Nielsen, R. ANGSD: analysis of next generation sequencing data. BMC Bioinformatics 15, 356 (2014).

[28] Rasmussen, M. et al. An Aboriginal Australian genome reveals separate human dispersals into Asia. Science 334, 94–98 (2011).

[29] Renaud, G., Slon, V., Duggan, A. T. & Kelso, J. Schmutzi: estimation of contamination and endogenous mitochondrial consensus calling for ancient DNA. Genome Biol. 16, 224 (2015).

[30] Rubinacci, S., Hofmeister, R. J., Sousa da Mota, B. & Delaneau, O. Imputation of low-coverage sequencing data from 150,119 UK Biobank genomes. Nat. Genet. 55, 1088–1090 (2023).

[31] Mallick, S. et al. The Allen Ancient DNA Resource (AADR): a curated compendium of ancient human genomes. Sci. Data 11, 182 (2024).

[32] Mallick, S. & Reich, D. The Allen Ancient DNA Resource (AADR): a curated compendium of ancient human genomes. Harvard Dataverse, V62.0 data release (16 September 2024), 10.7910/DVN/FFIDCW (2024).

[33] Fernandes, D. M., Cheronet, O., Gelabert, P. & Pinhasi, R. TKGWV2: an ancient DNA relatedness pipeline for ultra-low coverage whole genome shotgun data. Sci. Rep. 11, 21262 (2021).

[34] Andrews, R. M. et al. Reanalysis and revision of the Cambridge reference sequence for human mitochondrial DNA. Nat. Genet. 23, 147 (1999).

[35] Schönherr, S., Weissensteiner, H., Kronenberg, F. & Forer, L. Haplogrep 3—an interactive haplogroup classification and analysis platform. Nucleic Acids Res. 51, W263–W268 (2023).

[36] Weissensteiner, H. et al. HaploGrep 2: mitochondrial haplogroup classification in the era of high-throughput sequencing. Nucleic Acids Res. 44, W58–W63 (2016).

[37] Dür, A., Huber, N. & Parson, W. Fine-tuning phylogenetic alignment and haplogrouping of mtDNA sequences. Int. J. Mol. Sci. 22, 5747 (2021).

[38] Price, A. L. et al. Principal components analysis corrects for stratification in genome-wide association studies. Nat. Genet. 38, 904–909 (2006).

[39] Patterson, N., Price, A. L. & Reich, D. Population structure and eigenanalysis. PLoS Genet. 2, e190 (2006).

[40] Bergström, A. et al. Insights into human genetic variation and population history from 929 diverse genomes. Science 367, eaay5012 (2020).

[41] Mallick, S. et al. The Simons Genome Diversity Project: 300 genomes from 142 diverse populations. Nature 538, 201–206 (2016).

[42] Schiffels, S. PileupCaller: a command-line tool to sample genotypes from low-coverage sequencing data of ancient DNA. J. Open Source Softw. 11, 8634 (2026).

