## Supplementary Information for "Matched full-UDG and non-UDG ancient DNA libraries reveal trade-offs in post-mortem damage correction for imputation and kinship inference"

This file contains Supplementary Figures S1–S8 and Supplementary Tables S1–S10.

The publication-associated reproducibility release described under Code availability in the main manuscript contains the custom scripts, processed source data, workflow documentation, summary outputs, verification checksums and validation records supporting the main figures, tables and reported computational results. The exact original random state used during the primary READv2 pseudo-haploid sampling step was not preserved. The release provides anonymized pairwise $P_0$ tables, focal outputs, downstream commands, the normalisation procedure and validation records, allowing the published READv2 calculations to be verified. Full cemetery genotype matrices and controlled BAM inputs are not distributed.

mapDamage misincorporation profiles

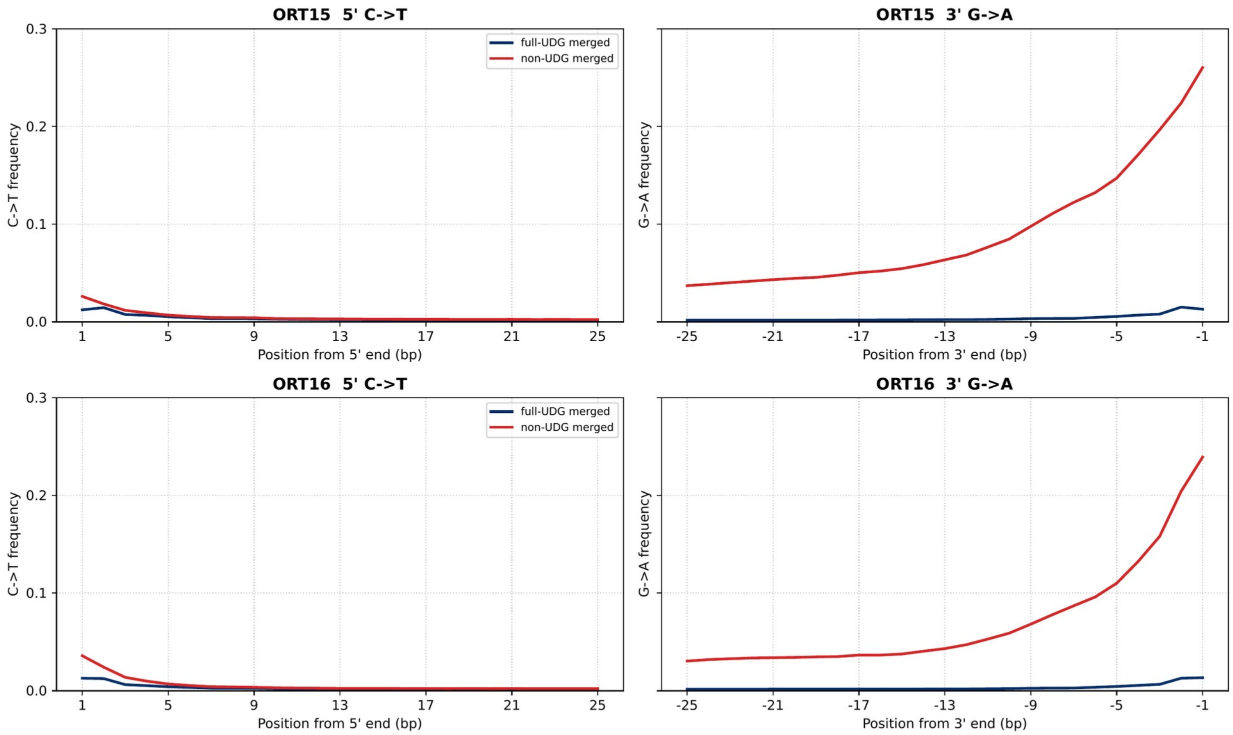


**Supplementary Figure S1. Post-mortem damage profiles for full-UDG and non-UDG libraries before computational correction.** mapDamage2 plots show 5′ C→T (left) and 3′ G→A (right) frequencies for ORT15 (top) and ORT16 (bottom). Full-UDG data are blue and show low 3′ G→A frequencies at position 1 (1.30% for ORT15 and 1.34% for ORT16); non-UDG data are red and reach 26.02% and 23.92%, respectively. Non-UDG 5′ C→T frequencies are also low (2.60% and 3.59%) and may reflect exposure to uracil-specific excision reagent (USER) during loop-adaptor cleavage. Profiles were calculated from merged collapsed-read BAMs before duplicate removal.

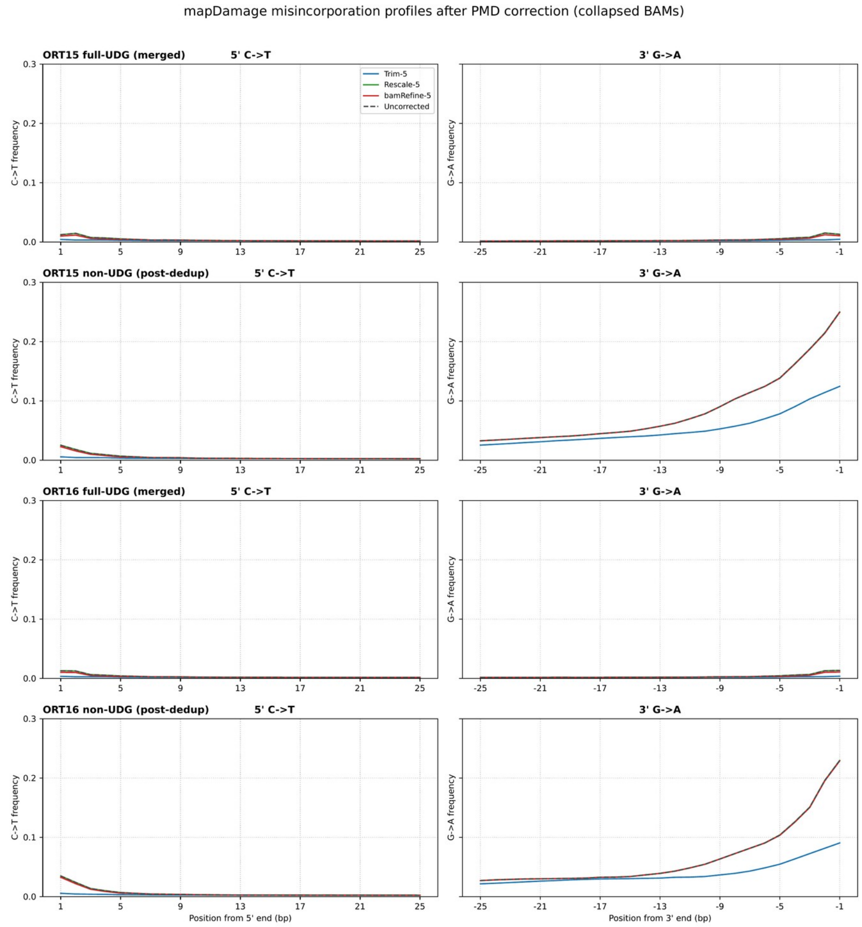


**Supplementary Figure S2. mapDamage2 misincorporation profiles before and after PMD correction.** The panels show 5′ C→T (left) and 3′ G→A (right) frequencies in uncorrected data and after 5-bp soft clipping, 5-bp mapDamage2 rescaling or 5-bp bamRefine masking. Full-UDG rows were calculated from merged collapsed-read BAMs before duplicate removal, whereas non-UDG rows use BAMs after duplicate removal, as labelled. All y axes span 0 to 30%. mapDamage2 counts observed substitutions without weighting revised base qualities, and bamRefine masks only bases at selected SNPs; their curves therefore need not flatten after correction. Soft clipping removes terminal bases and consequently flattens the curves near read ends. Because the library types were profiled at different processing stages, compare correction states within each row rather than quantitatively between full-UDG and non-UDG rows.

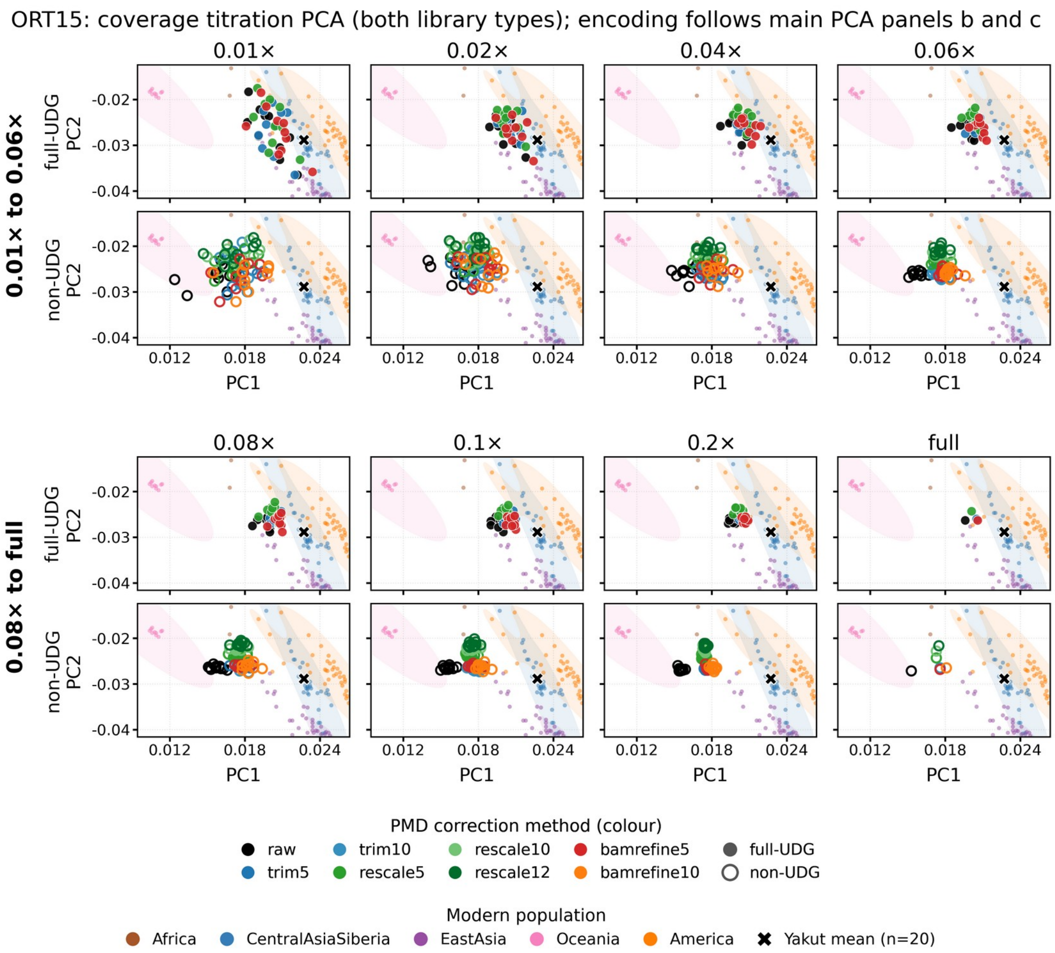


**Supplementary Figure S3. Coverage titration PCA for ORT15.** Full-UDG and non-UDG libraries were downsampled to 0.01–0.2×, with 10 replicates per attainable target, and analysed with the undownsampled source data. Genotypes were called pseudo-haploid and projected onto the AADR v62 HGDP+SGDP reference. The panels show 0.01× to 0.06× in the upper section and 0.08× to full in the lower section, with full-UDG above non-UDG. “Full” denotes the undownsampled source BAM for each correction state, not a common coverage; the corresponding depths are listed in Supplementary Table S10. Colours and symbols follow Figure 7, but this titration comes from a separate pseudo-haploid smartpca run. Scatter generally decreases with coverage, and full-UDG states form tighter clusters. Uncorrected non-UDG points are shifted relative to the matched full-UDG comparator, whereas corrected points are generally closer; these coordinates do not establish ancestry accuracy. ORT15 non-UDG Trim-10, with a source depth of about 0.17×, could not reach 0.2×.

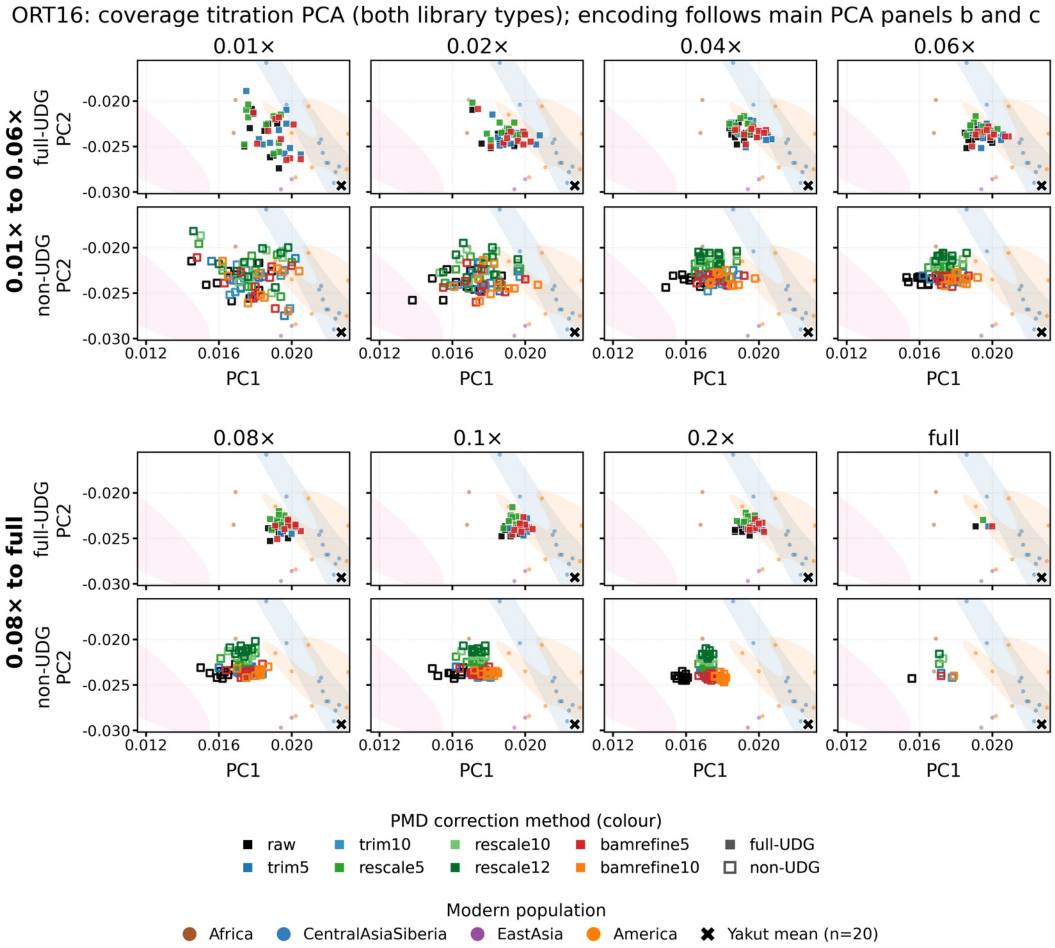


**Supplementary Figure S4. Coverage titration PCA for ORT16.** The design, coverage sections and encoding are as in Supplementary Figure S3. Pink shading represents the Oceania reference density; no Oceania reference points fall within the displayed range. “Full” again denotes the undownsampled source BAM for each correction state, with corresponding depths in Supplementary Table S10. All ORT16 correction states reached every target coverage. Scatter generally decreases with coverage, and full-UDG states form tighter clusters. Uncorrected non-UDG points are shifted relative to the matched full-UDG comparator, whereas corrected points are generally closer. These coordinates do not establish ancestry accuracy.

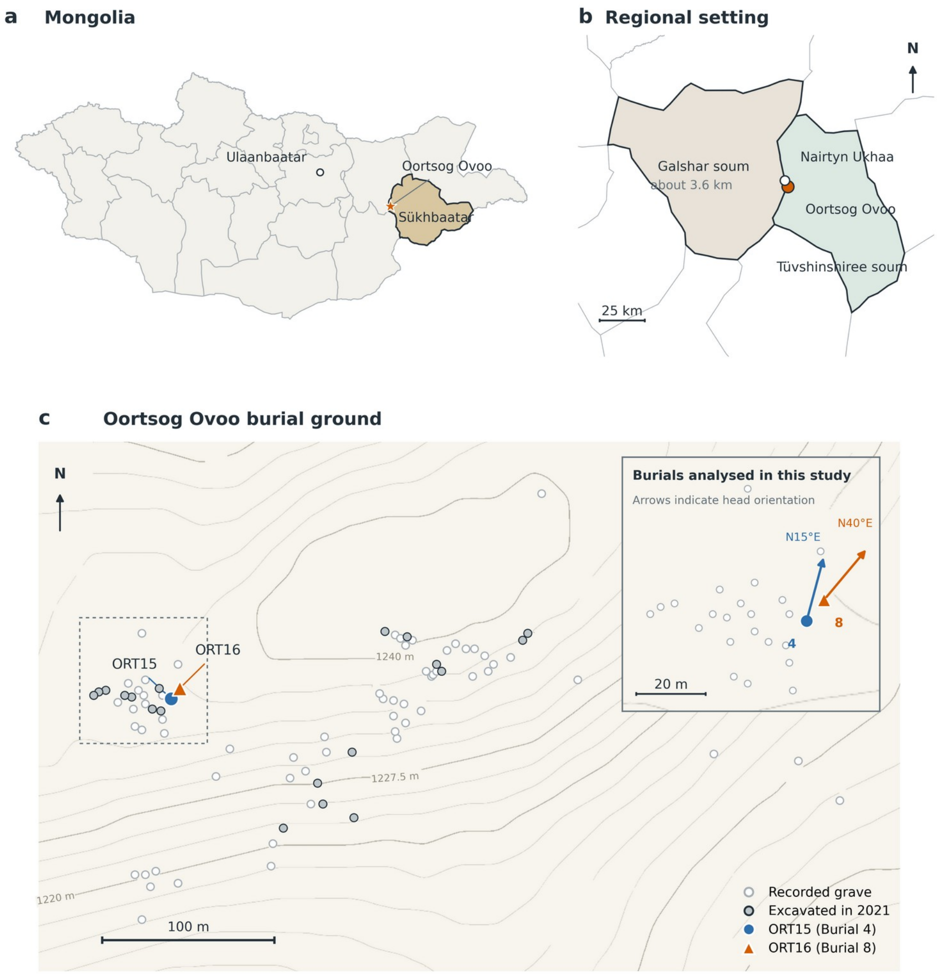


**Supplementary Figure S5. Geographic location and site plan of the Oortsog Ovoo burial ground.** (a) Location of Oortsog Ovoo in Mongolia, with Sükhbaatar Province highlighted. (b) Regional map showing Oortsog Ovoo and Nairtyn Ukhaa, about 3.6 km apart, near the boundary between Tüvshinshiree and Galshar soums. (c) Plan drawn from recorded coordinates showing 83 graves and topographic contours labelled in metres. Grey symbols denote graves excavated in 2021. ORT15 (Burial 4) and ORT16 (Burial 8) are highlighted, and the inset identifies the burials analysed in this study; arrows indicate head orientation (N15°E and N40°E). The plan and contours were redrawn from the 2022 report of the 2021 excavation cited in the main text; the original site plan was prepared by J. Erdene. Administrative boundaries are from geoBoundaries gbOpen MNG ADM1/ADM2, also cited in the main text (https://www.geoboundaries.org). Map lines delineate study areas and do not necessarily depict accepted national boundaries. Regional positions are approximate; exact archaeological coordinates are not displayed.

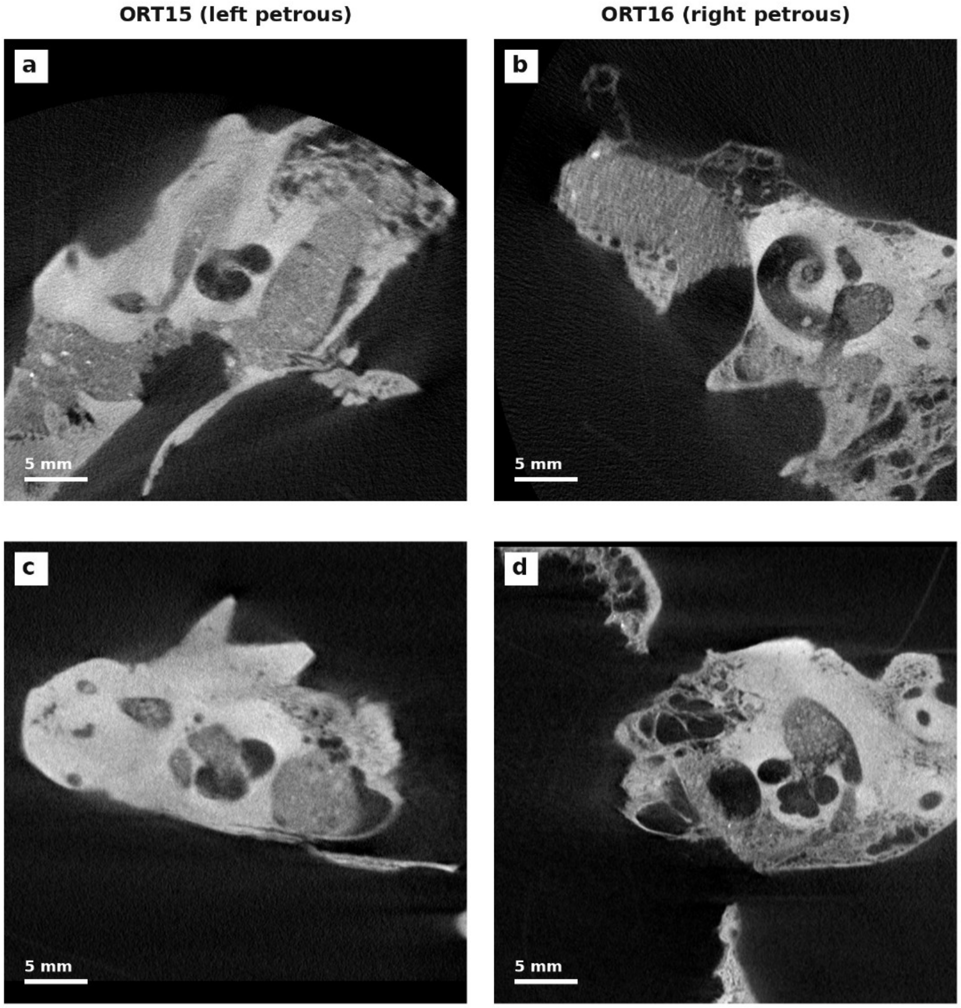


**Supplementary Figure S6. Pre-sampling micro-CT documentation of the petrous bones used for ancient DNA extraction.** Panels (a) and (c) show ORT15 (left petrous bone), and panels (b) and (d) show ORT16 (right petrous bone); each specimen is shown in two reconstructed views with identical display settings. Anatomical plane names are not assigned because orientation vectors were not preserved. Scanning preceded destructive sampling. Laboratory records identify the cochlear region as the sampling location and record powder masses of 26.5 mg for ORT15 and 36.2 mg for ORT16, but no coordinate map links these displayed sections to the exact drill position. No sampling arrows or drill-path annotations are therefore shown. Scanning used a Rigaku micro-CT system (90 kVp, 88 μA, Standard mode; 18 s scan, 72 mm field of view, Cu 0.06 + Al 0.5 filter), reconstructed to 1024×1024×1024 volumes with 72-μm isotropic voxels; scale bars, 5 mm. Images were used for anatomical localisation and documentation before sampling only; no quantitative CT analysis was performed.

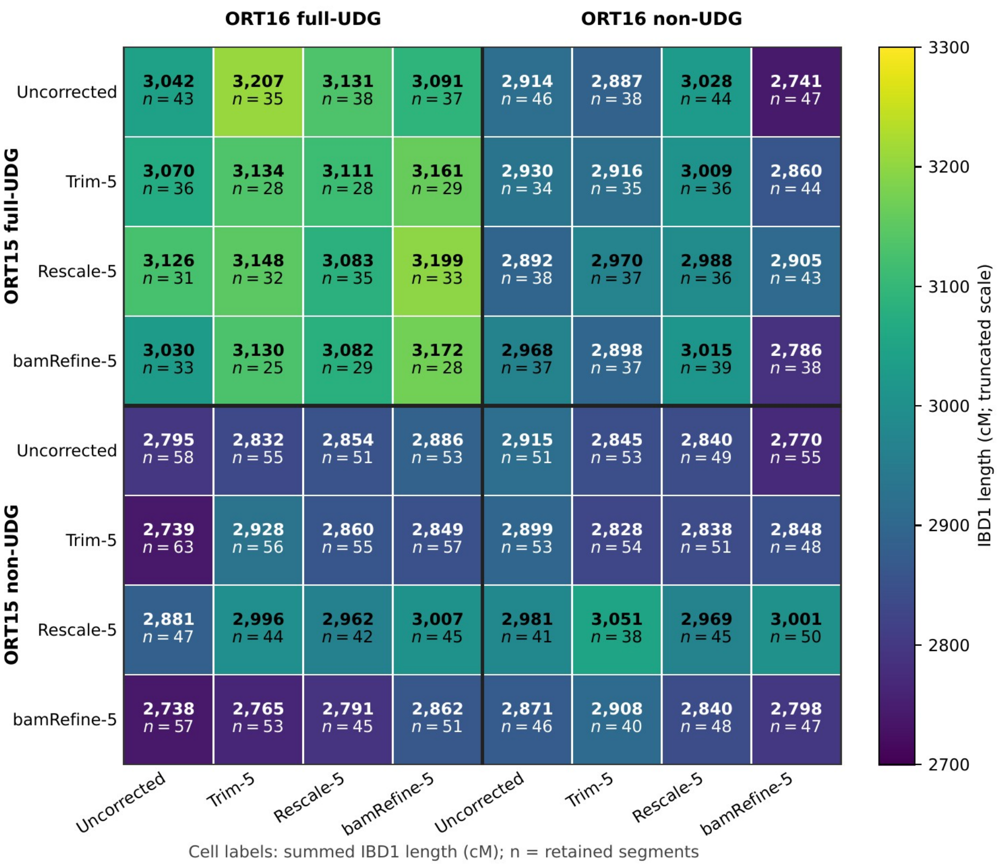


**Supplementary Figure S7. IBD1 comparison matrix across correction states.** The 8 × 8 matrix shows all 64 combinations of the eight ORT15 states (rows) and eight ORT16 states (columns). On each axis, the four full-UDG states precede the four non-UDG states. Each cell shows the summed length of IBD1 segments longer than 12 cM with marker density above 220 SNPs/cM; cM totals are rounded to the nearest whole number, and n is the number of retained segments. Thick lines separate the full-UDG and non-UDG blocks. Colour encodes summed IBD1 length on a truncated 2700–3300 cM scale. IBD1 was called with ancIBD 0.7 on the AADR v62 site set across a callable span of 3538.85 cM, without additional gap merging. Cells mirrored across the diagonal are not duplicates: they reverse which correction state is assigned to ORT15 and ORT16. A larger inferred IBD1 total does not by itself indicate greater accuracy.

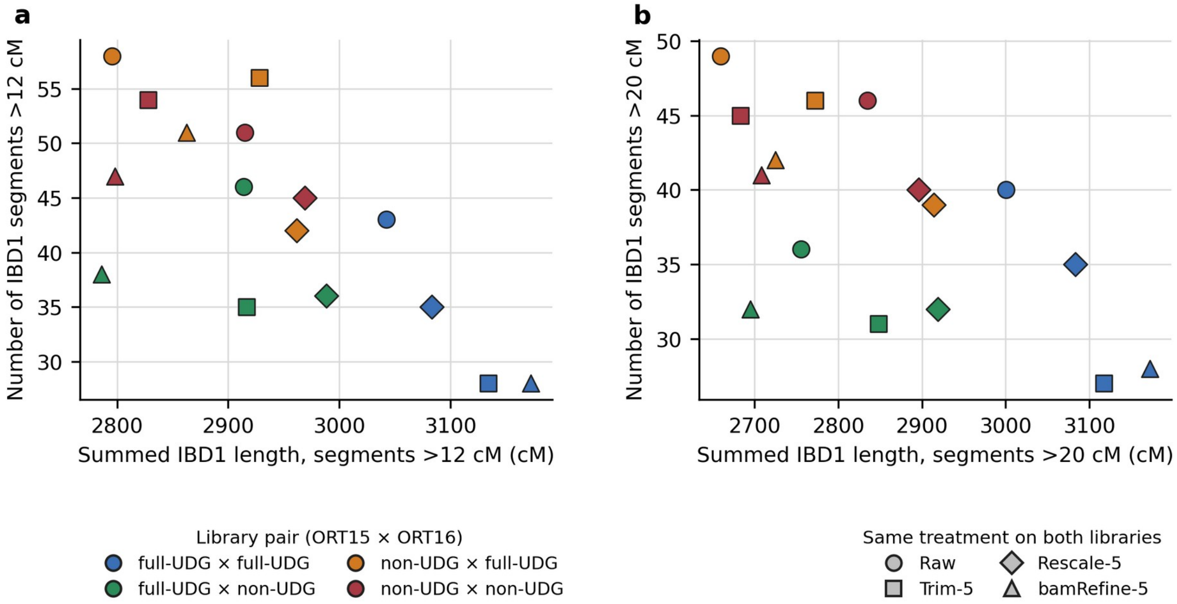


**Supplementary Figure S8. Summed IBD1 length and segment number across same-state library pairs.** Each point represents one of 16 combinations: four ORT15 and ORT16 library-type pairings, each evaluated with the same uncorrected, Trim-5, Rescale-5 or bamRefine-5 state applied to both libraries. Panels plot summed IBD1 length against the number of retained segments after requiring segments longer than (a) 12 cM or (b) 20 cM; all retained segments also had marker density above 220 SNPs/cM. Greater summed length indicates more inferred sharing, whereas a greater number of segments can reflect fragmentation. Neither measure alone ranks accuracy.

**Supplementary Table S1.** Per-library QC metrics. Endogenous DNA (%) was estimated with Qualimap on the pre-deduplication collapsed BAM, duplication rate (%) was calculated with samtools markdup, and mean fragment length was obtained from AdapterRemoval. The 3′ G→A value at position 1 is the terminal-base misincorporation frequency from the mapDamage2 plot on the merged pre-deduplication collapsed BAM (one value per individual per library type).

| Individual | Library | Endo. (%) | Dup. (%) | Length (bp) | 3′ G→A (%) |
|---|---|---|---|---|---|
| ORT15 | full-UDG library a | 85.1 | 64.9 | 60.6 | — |
| | full-UDG library b | 83.5 | 56.8 | 61.7 | — |
| | full-UDG library c | 82.1 | 56.8 | 62.0 | — |
| | full-UDG merged (a+b+c) | — | — | — | 1.30 |
| | non-UDG | 81.5 | 40.8 | 63.4 | 26.02 |
| ORT16 | full-UDG library a | 90.8 | 44.5 | 59.1 | — |
| | full-UDG library b | 88.9 | 38.5 | 64.1 | — |
| | full-UDG library c | 88.7 | 56.0 | n/a‡ | — |
| | full-UDG merged (a+b+c) | — | — | — | 1.34 |
| | non-UDG | 89.3 | 41.3 | 64.4 | 23.92 |

The full-UDG 3′ G→A values were measured on the merged a+b+c data; individual reactions were not assayed separately. ‡ No fragment-length result was available.

**Supplementary Table S2. Present-day contamination estimates for ORT15 and ORT16.** Nuclear contamination was estimated from the X chromosome with ANGSD v0.933 (Method 1) on the collapsed full-UDG BAM using the HapMapChrX panel; mitochondrial contamination was estimated with the standalone mtCont component of schmutzi v1.5.7 on the collapsed, deduplicated non-UDG BAM. Nuclear estimates are given with their standard errors; mtCont uncertainties are the program-reported 95% intervals. The mitochondrial minor-allele quality-control statistic is not a contamination estimator and is not tabulated here.

| Individual | Genome | Input library | Method | Estimate | Uncertainty | Informative sites |
|---|---|---|---|---|---|---|
| ORT15 | Nuclear X† | Full-UDG | ANGSD Method 1, MoM§ | 0.22% | SE 0.08% | 6757 |
| ORT15 | Nuclear X† | Full-UDG | ANGSD Method 1, ML§ | 0.39% | SE 0.12% | 6757 |
| ORT15 | mtDNA | Non-UDG | Standalone mtCont‡ | 1% | 0–2% | — |
| ORT16 | mtDNA | Non-UDG | Standalone mtCont‡ | 1% | 0–2% | — |

† Nuclear X-chromosome estimation was applicable only to genetically male ORT15.

‡ mtCont was run as a standalone analysis, not as the complete iterative schmutzi workflow.

§ MoM denotes method of moments and ML denotes maximum likelihood. ANGSD Method 2 produced concordant values (MoM 0.26%, ML 0.30%), retained in the publication-associated reproducibility repository release described on page 1 rather than presented as additional primary estimates.

**Supplementary Table S3.** Alternate-read fractions and covered-site retention after PMD correction, including a targeted 3′-end-only trimming sensitivity analysis. Panel A reports correction-state summaries. Trim-5 and Trim-10 soft-clip the stated number of bases from both read ends; 3′-only-5 and 3′-only-10 soft-clip the stated number only from the strand-aware 3′ end. Means were calculated separately at damage-prone C→T/G→A SNPs and transversion (Tv) SNPs with combined REF+ALT depth of at least three reads. ΔALT is the damage-prone mean minus the transversion mean. Covered sites had combined REF+ALT depth of at least one read, and retention is relative to the uncorrected library of the same individual and library type. Transition and transversion classes may differ in allele-frequency and reference/alternate composition, and callable sites remain method-specific; ΔALT is therefore descriptive, and lower values do not by themselves demonstrate loss of genuine variants. ΔALT entries give the pooled point estimate followed by the leave-one-autosome-out 95% confidence interval. Prespecified paired-contrast conclusions under the correction-specific primary analysis were unchanged in fixed 5-Mb and 10-Mb block sensitivity analyses. Full primary and common-callable paired-contrast results are provided in the accompanying Supplementary Data.

**A. Correction-state summaries.**

| Library | Strategy | Damage-prone (%) | Tv (%) | ΔALT (pp; 95% CI) | Covered (M) | Retained (%) |
|---|---|---|---|---|---|---|
| **Full-UDG** | | | | | | |
| ORT15 | Uncorrected | 3.53 | 3.06 | +0.47 (+0.41 to +0.54) | 28.85 | 100.0 |
| | Trim-5 | 2.93 | 3.01 | −0.09 (−0.16 to −0.01) | 24.34 | 84.4 |
| | Rescale-5 | 2.23 | 2.99 | −0.76 (−0.82 to −0.69) | 28.74 | 99.6 |
| | Rescale-12 | 1.23 | 2.96 | −1.73 (−1.79 to −1.66) | 28.66 | 99.3 |
| | bamRefine-5 | 2.94 | 3.00 | −0.06 (−0.13 to +0.01) | 26.62 | 92.3 |
| ORT16 | Uncorrected | 3.37 | 3.03 | +0.34 (+0.29 to +0.39) | 45.57 | 100.0 |
| | Trim-5 | 2.84 | 2.98 | −0.13 (−0.18 to −0.08) | 40.12 | 88.0 |
| | Rescale-5 | 2.22 | 2.97 | −0.74 (−0.79 to −0.70) | 45.45 | 99.7 |
| | Rescale-12 | 1.32 | 2.95 | −1.63 (−1.69 to −1.58) | 45.35 | 99.5 |
| | bamRefine-5 | 2.86 | 2.98 | −0.11 (−0.16 to −0.06) | 42.88 | 94.1 |
| **Non-UDG** | | | | | | |
| ORT15 | Uncorrected | 4.33 | 3.48 | +0.85 (+0.74 to +0.97) | 14.38 | 100.0 |
| | Trim-5 | 3.67 | 3.26 | +0.41 (+0.27 to +0.57) | 12.18 | 84.7 |
| | 3′-only-5 | 3.78 | 3.23 | +0.55 (+0.41 to +0.69) | 13.44 | 93.4 |
| | Trim-10 | 3.43 | 3.17 | +0.26 (+0.06 to +0.47) | 9.92 | 69.0 |
| | 3′-only-10 | 3.57 | 3.11 | +0.46 (+0.32 to +0.61) | 12.49 | 86.9 |
| | Rescale-5 | 2.92 | 3.17 | −0.25 (−0.38 to −0.12) | 14.27 | 99.3 |
| | Rescale-10 | 2.03 | 3.04 | −1.01 (−1.12 to −0.89) | 14.21 | 98.8 |
| | Rescale-12 | 1.75 | 3.00 | −1.24 (−1.35 to −1.13) | 14.20 | 98.7 |
| | bamRefine-5 | 3.60 | 3.19 | +0.41 (+0.28 to +0.55) | 13.26 | 92.2 |
| | bamRefine-10 | 3.31 | 3.04 | +0.27 (+0.13 to +0.42) | 12.12 | 84.3 |

| Library | Strategy | Damage-prone (%) | Tv (%) | ΔALT (pp; 95% CI) | Covered (M) | Retained (%) |
|---|---|---|---|---|---|---|
| ORT16 | Uncorrected | 4.08 | 3.44 | +0.64 (+0.52 to +0.76) | 17.63 | 100.0 |
| | Trim-5 | 3.40 | 3.17 | +0.23 (+0.09 to +0.38) | 15.05 | 85.3 |
| | 3′-only-5 | 3.50 | 3.18 | +0.32 (+0.18 to +0.46) | 16.55 | 93.9 |
| | Trim-10 | 3.26 | 3.13 | +0.14 (−0.05 to +0.32) | 12.36 | 70.1 |
| | 3′-only-10 | 3.35 | 3.11 | +0.24 (+0.07 to +0.41) | 15.48 | 87.8 |
| | Rescale-5 | 2.71 | 3.14 | −0.43 (−0.54 to −0.32) | 17.51 | 99.3 |
| | Rescale-10 | 1.92 | 3.05 | −1.12 (−1.23 to −1.02) | 17.45 | 99.0 |
| | Rescale-12 | 1.68 | 3.02 | −1.35 (−1.45 to −1.25) | 17.44 | 98.9 |
| | bamRefine-5 | 3.39 | 3.14 | +0.25 (+0.12 to +0.38) | 16.29 | 92.4 |
| | bamRefine-10 | 3.16 | 3.05 | +0.11 (−0.03 to +0.25) | 14.95 | 84.8 |

**B. Paired 3′-only versus symmetric Trim-5 contrasts.**

| Individual | Contrast | Retention difference (percentage points) | Observed ΔALT difference (percentage points) | Bias-corrected ΔALT difference (percentage points) | Jackknife SE | 95% CI for ΔALT difference (percentage points) |
|---|---|---|---|---|---|---|
| ORT15 | 3′-only-5 − Trim-5 | +8.7159 | +0.1329 | +0.1330 | 0.0286 | +0.0735 to +0.1925 |
| ORT15 | 3′-only-10 − Trim-5 | +2.1593 | +0.0468 | +0.0460 | 0.0443 | −0.0462 to +0.1382 |
| ORT16 | 3′-only-5 − Trim-5 | +8.5534 | +0.0884 | +0.0882 | 0.0147 | +0.0576 to +0.1188 |
| ORT16 | 3′-only-10 − Trim-5 | +2.4413 | +0.0097 | +0.0084 | 0.0259 | −0.0455 to +0.0623 |

Differences are calculated as the 3′-only state minus symmetric Trim-5. The 3′-only-5 comparison is nominal-window matched; 3′-only-10 versus Trim-5 removes the same total number of terminal bases. Positive retention differences indicate greater covered-site retention under 3′-only trimming, whereas positive differences in ΔALT indicate more residual damage-associated alternative-allele support. Paired leave-one-autosome-out jackknifing omitted the same autosome from both correction-specific state aggregates. Sites were not required to be identical, so pairing is by omitted autosome rather than individual site. Confidence intervals use the bias-corrected jackknife estimate; fixed 5-Mb and 10-Mb block analyses gave the same inference.

**Supplementary Table S4.** BamRefine-5 versus Rescale-5 non-reference discordance and paired delete-one-autosome jackknife results.

| Individual | bamRefine-5 NRD (%) | Rescale-5 NRD (%) | Observed ΔNRD (percentage points) | Bias-corrected ΔNRD (percentage points) | Jackknife SE (percentage points) | 95% CI (percentage points) |
|---|---:|---:|---:|---:|---:|---|
| ORT15 | 18.2961 | 18.3164 | −0.0203 | −0.0208 | 0.0875 | −0.2028 to +0.1612 |
| ORT16 | 14.8823 | 14.8521 | +0.0302 | +0.0302 | 0.0660 | −0.1072 to +0.1675 |

ΔNRD was defined as bamRefine-5 NRD minus Rescale-5 NRD. For each individual, the two method-specific NRDs were calculated independently against the same fixed full-UDG comparator, reference-frequency panel, filters and allele-frequency bins. Uncertainty was estimated with a paired delete-one-autosome jackknife, omitting the same autosome from both method-specific count aggregates in each replicate. Site sets were not constrained to be identical between methods; pairing therefore operated at the level of omitted chromosomes rather than individual sites. The 95% intervals use the bias-corrected jackknife estimate, jackknife standard error and the t distribution with 21 degrees of freedom. NRD, ΔNRD, SE and interval values are expressed in percentage points where indicated.

**Supplementary Table S5.** Inferred IBD1 for the asymmetric direct-pair comparison. One individual's full-UDG library remains uncorrected while only the other's non-UDG correction varies, in both directions. Columns give summed IBD1 length at strict >8 and >12 cM thresholds and segment counts at strict >12 and >20 cM; all also require marker density >220 SNP/cM. These values measure agreement between library preparations, not genotype accuracy; a larger total is not by itself evidence of greater accuracy.

| Strategy (non-UDG side) | IBD1 >8 cM (cM) | IBD1 >12 cM (cM) | N >12 | N >20 |
|---|---|---|---|---|
| **ORT15 full-UDG (uncorrected) × ORT16 non-UDG** | | | | |
| Uncorrected | 2922.1 | 2913.9 | 46 | 36 |
| Trim-5 | 2919.7 | 2887.1 | 38 | 35 |
| Rescale-5 | 3037.6 | 3027.9 | 44 | 37 |
| bamRefine-5 | 2773.5 | 2741.2 | 47 | 39 |
| **ORT15 non-UDG × ORT16 full-UDG (uncorrected)** | | | | |
| Uncorrected | 2857.3 | 2795.5 | 58 | 49 |
| Trim-5 | 2804.2 | 2739.3 | 63 | 50 |
| Rescale-5 | 2900.4 | 2880.5 | 47 | 41 |
| bamRefine-5 | 2771.7 | 2738.2 | 57 | 45 |

**Supplementary Table S6.** TKGWV2 treatment-series estimates and READv2 results for ORT15 and ORT16. For TKGWV2, the coefficient is HRC; values above 0.1875 are first-degree, whereas values from 0.0625 through 0.1875 are second-degree. Directional mixed-library rows identify the library type assigned to each individual. The two READv2 rows report the primary 1240k analysis and an autosome-only 1240k rerun; their normalisation cohorts contained 21 and 22 retained individuals, respectively. Only the ORT15–ORT16 result is shown. For READv2, the coefficient is KC. SNPs denotes used SNPs for TKGWV2 and overlapping SNPs for READv2.

| Pairing | State | SNPs | Coefficient | Classification |
|---|---|---|---|---|
| Full-UDG × full-UDG | Uncorrected | 5,170,890 | 0.2096 | First-degree |
| | bamRefine-5 | 4,517,214 | 0.2356 | First-degree |
| | Rescale-5 | 5,110,222 | 0.2444 | First-degree |
| | Trim-5 | 3,880,873 | 0.2333 | First-degree |
| Non-UDG × non-UDG | Uncorrected | 981,982 | 0.1417 | Second-degree |
| | bamRefine-5 | 843,168 | 0.1854 | Second-degree |
| | Rescale-5 | 966,422 | 0.1931 | First-degree |
| | Trim-5 | 717,822 | 0.1765 | Second-degree |
| ORT15 non-UDG × ORT16 full-UDG | Uncorrected | 2,530,441 | 0.1680 | Second-degree |
| | bamRefine-5 | 2,209,919 | 0.2058 | First-degree |
| | Rescale-5 | 2,497,066 | 0.2131 | First-degree |
| | Trim-5 | 1,905,460 | 0.2008 | First-degree |
| ORT15 full-UDG × ORT16 non-UDG | Uncorrected | 1,977,690 | 0.1765 | Second-degree |
| | bamRefine-5 | 1,698,223 | 0.2122 | First-degree |
| | Rescale-5 | 1,949,002 | 0.2203 | First-degree |
| | Trim-5 | 1,436,949 | 0.2089 | First-degree |
| READv2: ORT15–ORT16 | Primary 1240k | 169,172 | 0.230967 | First-degree; parent–offspring |
| | Autosome-only 1240k | 157,451 | 0.231592 | First-degree; parent–offspring |

**Supplementary Table S7.** Mitochondrial haplogroup assignments and call-set concordance. Variants were called from the full-UDG collapsed-only .1 BAM with bcftools v1.16 using MAPQ ≥30, base quality ≥30, BAQ disabled and haploid calling; no global minimum-depth, allele-fraction or strand-balance filter was applied. Shared calls require identical chromosome, position, REF and ALT. The MT:3106 CN>C record is the left-anchored deletion of the artificial rCRS N spacer at position 3107 and was excluded from biological SNP and discordance counts. Absence of a call at zero qualifying depth was not interpreted as a reference genotype.

| Sample | Input | Haplogroup | Score | ALT SNPs | Exact shared | Additional call | Interpretation |
|---|---|---|---|---|---|---|---|
| ORT15 | Full-UDG collapsed-only .1 | D4o1 | 0.8911 | 36 | 36 | None | The ORT15 call set is an exact subset of the ORT16 caller-emitted SNP set. |
| ORT16 | Full-UDG collapsed-only .1 | D4o1 | 0.9107 | 37 | 36 | C7028T | One-read caller-emitted call; not a confirmed between-individual difference. |

The additional C7028T call in ORT16 was supported by one read; ORT15 had no qualifying coverage at this position, so a between-individual difference could not be established.

**Supplementary Table S8. Accelerator mass spectrometry (AMS) radiocarbon dating result for ORT16** (Paleo Labo Co., Ltd.; bone collagen). Calendar-age ranges are reproduced as reported by the laboratory. The calibration software and curve were not specified. ORT15 was not radiocarbon dated.

| Field | ORT16 result |
|---|---|
| Laboratory code | PLD-55920 |
| $\delta^{13}C$ | −18.10 ± 0.19‰ |
| $^{14}C$ age | 705 ± 15 yr BP |
| Fractionation-corrected age used for calibration | 707 ± 17 yr BP |
| 1σ range | 1278–1292 cal AD (68.27%) |
| 2σ ranges | 1274–1299 cal AD (92.62%); 1372–1376 cal AD (2.83%) |

**Supplementary Table S9.** Core software and external-resource versions and provider sources. Recorded parameters and portable workflow scripts are provided in the publication-associated reproducibility release described under Code availability in the main manuscript. "Not recorded" is used where the archived environment did not preserve a more specific build identifier.

| Software or resource | Version/build used | Project/provider source |
|---|---|---|
| AdapterRemoval | 2.2.2 | https://github.com/MikkelSchubert/adapterremoval |
| BWA | 0.7.17-r1188 | https://github.com/lh3/bwa |
| samtools | 1.7 | https://www.htslib.org |
| Qualimap | 2; exact build not recorded | http://qualimap.conesalab.org |
| bcftools/HTSlib | 1.16/1.16 | https://www.htslib.org |
| trimBam (BamUtil) | 1.0.15 | https://genome.sph.umich.edu/wiki/BamUtil |
| mapDamage2 | 2.3.0a0-5175c20 | https://github.com/ginolhac/mapDamage |
| bamRefine | 0.2.1 | https://github.com/etkayapar/bamRefine |
| GLIMPSE | 1.1.1 | https://odelaneau.github.io/GLIMPSE |
| GLIMPSE2 concordance | 2.0.0; commit 3bed6d9 | https://odelaneau.github.io/GLIMPSE |
| ancIBD | 0.7 | https://github.com/hringbauer/ancIBD |
| ANGSD | 0.933 | https://github.com/ANGSD/angsd |
| schmutzi | 1.5.7 | https://github.com/grenaud/schmutzi |
| TKGWV2 | 1.0b | https://github.com/danimfernandes/tkgwv2 |
| READv2 | 2.01 | https://github.com/GuntherLab/READv2 |
| PLINK | 1.90b6.24 | https://www.cog-genomics.org/plink |
| PLINK 2 | 2.0.0-a.6.4LM, 6 Dec 2024 | https://www.cog-genomics.org/plink/2.0 |
| pileupCaller (sequenceTools) | 1.5.4.0 | https://github.com/stschiff/sequenceTools |
| HaploGrep 3 | 3.3.2 | https://haplogrep.i-med.ac.at |
| PhyloTree 17 Forensic Update | rCRS tree 1.3 | https://haplogrep.i-med.ac.at/phylogenies |
| smartpca (EIGENSOFT) | build 18140 | https://github.com/DReichLab/EIG |
| hs37d5 reference | GRCh37/hs37d5; FASTA SHA-256 65add55817fc9a5de8221caf36e84bf867 0f94bef9cff7889093692b271a765d | https://ftp.1000genomes.ebi.ac.uk/vol1/ftp/technical/reference/ phase2_reference_assembly_sequence/ |
| 1000 Genomes reference and SNP panels | Phase 3, GRCh37; 2504-sample SHAPEIT2 integrated v5b panel and matched biallelic sites | https://www.internationalgenome.org/data-portal/data-collection/phase-3 |
| Genetic maps | GRCh37 maps accompanying the Phase 3 SHAPEIT2 integrated v5b panel | https://odelaneau.github.io/GLIMPSE/ |
| Allen Ancient DNA Resource | AADR v62.0, 1240k panel | https://reich.hms.harvard.edu/allen-ancient-dna-resource-aadr-downloadable-genotypes-present-day-and-ancient-dna-data |

**Supplementary Table S10. Undownsampled source depths for coverage titration PCA.** Depths used for downsampling were calculated across autosomes 1–22 with samtools coverage. They differ slightly from the rounded Qualimap coverages shown in Figure 1 of the main text because the two summaries use different coverage calculations. States grouped in one row used the same undownsampled source depth. These values define the “full” column in Supplementary Figures S3 and S4; “full” does not denote a common depth.

| Individual | Library type | State or states | Source depth (×) |
|---|---|---|---|
| ORT15 | full-UDG | Uncorrected, Rescale-5 and bamRefine-5 | 0.863 |
| ORT15 | full-UDG | Trim-5 | 0.694 |
| ORT15 | non-UDG | Uncorrected, Rescale-5, Rescale-10, Rescale-12, bamRefine-5 and bamRefine-10 | 0.263 |
| ORT15 | non-UDG | Trim-5 | 0.218 |
| ORT15 | non-UDG | Trim-10 | 0.173 |
| ORT16 | full-UDG | Uncorrected, Rescale-5 and bamRefine-5 | 1.411 |
| ORT16 | full-UDG | Trim-5 | 1.144 |
| ORT16 | non-UDG | Uncorrected, Rescale-5, Rescale-10, Rescale-12, bamRefine-5 and bamRefine-10 | 0.339 |
| ORT16 | non-UDG | Trim-5 | 0.281 |
| ORT16 | non-UDG | Trim-10 | 0.224 |
